# Macronutrient Distribution and Protein Secondary Structure of Cool-Season Oats Revealed by Synchrotron-Based Mid-IR Spectroscopy and FTIR Chemical Imaging: Effects of Variety and Steam-Pressure Toasting Duration

**DOI:** 10.64898/2026.08.20.746060

**Authors:** Ganqi Deng, Maria E. Rodriguez-Espinosa, Kaiyang Tu, Jarvis Stobbs, Miranda Vu, Chithra Karunakaran, Xin Feng, Fang-Xiang Wu, Peiqiang Yu

## Abstract

This study aims to investigate changes in protein secondary structures (α-helix, 0-sheet, random coils, and *β*-turn) and macronutrient distribution in different cool-season oat varieties and steam­pressure toasting durations using synchrotron-based mid-infrared (Mid-IR) spectroscopy and Fourier Transform Infrared spectroscopy (FTIR) imaging. All oat samples, provided by the Crop Development Center at the University of Saskatchewan, were harvested over three consecutive years (2018, 2019, and 2020). The first experiment compared four oat varieties (CDC Arborg, CDC Nasser, CDC Haymaker, and Summit), while the second examined CDC Nasser oats subjected to steam-pressure toasting (SPT) at 121 for 0, 30, 60, 90, and 120 minutes. FTIR chemical imaging revealed that carbohydrates, proteins and lipids in the four oat varieties were mainly concentrated in the endosperm, aleurone layer and embryo, crease region, and remained unchanged after SPT. Peak-fitting deconvolution of the Amide I band (1700-1600 cm^-1^) and subsequent quantitative analysis revealed that the four oat varieties exhibited broadly similar protein secondary structure profiles, with statistically significant but subtle variety effects detected for *a*-helix (*P = 0.026*), 0-turn (*P = 0.047*), and the *a*-helix to *β*-sheet ratio (*P = 0.048*), however, *β*-sheet and random coil proportions did not differ significantly among varieties. In contrast, SPT induced pronounced structural rearrangements, with significant increase in *β*-sheet proportion (*P = 0.003*) and significant decreases in random coil content (*P = 0.026*). Notably, 30 minutes of toasting was sufficient to significantly increase *β*-sheet and decrease the random coil contents. These changes are consistent with heat-induced protein denaturation and intermolecular *β*-sheet aggregation, where thermal energy breaks the hydrogen bonds that stabilize the disordered random coil conformation, causing the unfolded polypeptide chains to reassemble into highly ordered *β*-sheet aggregates. After SPT, the peak centers of Amide I and II bands shifted to lower wavenumbers and both bands broadened while their intensities were maintained, reflecting the reorganization of the remaining protein into 0-sheet aggregates rather than any loss of amide-active protein. These findings suggest that, although genotype has a relatively minor effect on the protein secondary structure of oats, hydrothermal treatments fundamentally reorganize the protein matrix from a disordered to an ordered conformation, which may have implications for protein digestibility, solubility, and nutritional function.

## 1. Introduction

Oats (*Avena sativa L.*) grow primarily in cool and humid regions of the northern hemisphere with Canada representing one of the largest producers and exporters of oats (Statistics Canada, 2023). Compared with other cereals, oats have a much higher protein content and are rich in essential amino acids such as lysine and threonine (Klose and Arendt, 2012). Oats also have the highest oil content among the common cereals, reaching 18% in some varieties (Banaś et al., 2007). Oil is stored in oil bodies distributed throughout the endosperm, embryo and scutellum. In addition to the major macronutrients, oats also have higher levels of phytochemicals and other nutrients, such as fatty acids, 0-glucan and phenolic compounds than other cereals (Deng et al., 2023a; Tosh and Miller, 2016; Tosta et al., 2024). They are also a good source of magnesium, calcium, manganese, iron, copper, zinc, and selenium (Deng et al., 2025, 2023b). Many studies have shown that eating oats has numerous health benefits, such as reducing blood cholesterol and the incidence of coronary heart disease (CHD) (Hu et al., 2022; Li et al., 2024; Tosh and Bordenave, 2020), protecting against cancer (Li et al., 2022; Paudel et al., 2021), and preventing cardiovascular disease (Glenn et al., 2023; Llanaj et al., 2022; Wu et al., 2019).

The protein composition of oats differs significantly from that of wheat, barley and corn. While the storage proteins in these grains primarily consisted of alcohol-soluble proteins, the major storage proteins in oats are mainly the avenalin and salt-soluble 12S globulin, consisting of disulfide bonds linked by a- and *β*-subunits (Klose and Arendt, 2012; Peterson, 1978). The remaining protein components included albumin, glutelin, prolamin, enzymes and enzyme inhibitors. This molecular organization is highly beneficial for the structural stability of oat globulin in oats, but it may also limit its solubility and functional properties in the food system (Li et al., 2025). Oat protein functionality is therefore influenced not only by amino acid composition and concentration, but also by conformation, aggregation, and interaction with other components. The composition of oat grains varies greatly among different varieties and growing conditions. According to surveys and reports on various oat varieties, there is a wide variation in their protein, fat, starch, and 0-glucan concentrations. For example, the literature reported significant differences in protein concentration, ranging from 11.5 to 22.9% on a dry matter basis, among varieties grown under common conditions (Lampoglou et al., 2026). A study of thirty different oat varieties in Canada also found that protein, starch, and amylose contents were significantly influenced by varieties, environment, and their interactions (Alexander et al., 2025). Early studies have also shown that protein and *β*-glucan contents were affected by genetic and environmental factors, while fat concentration was strongly influenced by genotype (Doehlert et al., 2001). However, compositional analysis only revels how much protein a variety contains, rather than how that protein is organized at the molecular level. As a result, it remains unclear whether genotypic differences in overall composition are accompanied by systematic differences in the secondary structure of proteins within the whole grain tissue.

Hydrothermal treatment is another major factor affecting oat quality as oat grains contain high levels of endogenous lipase activity. This enzyme rapidly hydrolyzes triglycerides into free fatty acids and causes hydrolytic rancidity during storage. This lipase activity is inactivated by steam treatment, whereas the free fatty acid contents can exceed 30% of the fat phase within a few months of storage in oats that have not undergone proper heat treatment (Ekstrand et al., 1993). Thermal stabilization is therefore a critical step in commercial oat processing and is typically carried out in conjunction with dehulling, cutting, rolling, or grinding oats (Decker et al., 2014). Consequently, the protein matrix in most commercial oat products is already exposed to high temperature and moisture prior to further processing or consumption. Heat treatment can simultaneously alter the solubility, extractability, and functionality of protein while controlling enzymatic rancidity. Previous studies have shown that processing can change the infrared spectra of Amide I and Amide II in oat grains and their protein degradation characteristics (Tosta et al., 2020). The extent and rate of protein structural reorganization depend on the processing temperature, duration, moisture availability and physical state of the grain matrix. Steam-pressure toasting (SPT) exposes the whole grains to steam at a controlled temperature for a specified duration, providing a useful model for studying this response. SPT allows rapid structural changes to be distinguished from those that develop gradually over extended processing periods. Until now, most studies on the structure of oat protein have been conducted using extracted globulin fractions or isolated proteins. Although such systems are useful for characterizing intrinsic protein properties, the extraction process alters the aggregation state and molecular conformation. It also removes the proteins from the spatially heterogeneous environment of the grain, where they coexist with starch granules, lipids, cell wall polysaccharides and other macromolecules. Therefore, structural responses measured in protein isolates may differ from those occurring in intact oat tissue. Direct measurements within grain sections are needed to determine how protein conformation is organized across anatomical regions and how it responds to manipulation of its natural matrix.

Infrared spectroscopy is well-suited for this study because molecular functional groups can be examined without extensive chemical extraction. The Amide I region is sensitive to the protein secondary structures and is primarily located between 1700 and 1600 cm^-1^ and dominated by the stretching vibration of the C=O bond in the peptide backbone. Both resolution enhancement and peak fitting procedures can be used to estimate the relative contributions of *a*-helices, *β*-sheets, random coils and *β*-turns (Barth, 2007; Byler and Susi, 1986; Dong et al., 1990). However, these estimates are strongly dependent on spectral preprocessing, baseline correction, the number and form of fitted components, and the constraints applied during curve fitting. The protein secondary structure proportions obtained from infrared spectroscopy should be interpreted as comparative spectral estimates rather than absolute structural measurements (Surewicz et al., 1993; Yu, 2005). The analysis of intact grain is also limited by signal intensity and spatial resolution. Traditional globar-based infrared spectroscopy can image large tissue areas effectively, but its brightness becomes limiting when spectra are collected through micrometer-scale apertures. Synchrotron-based infrared spectroscopy overcomes this limitation, as the high brightness and small effective source size of synchrotron radiation allow the collection of spectra with a high signal-to-noise ratio near the diffraction limit (Miller and Dumas, 2006). Fourier Transform Infrared Spectroscopy (FTIR) imaging can visualize the distribution of protein, carbohydrate, and lipid-related signals across the entire cross-section of the seeds, while synchrotron-based point spectroscopy provides higher-quality spectra from selected anatomical regions for detailed structural analysis of protein, carbohydrates, and lipids.

Although interest in oat protein is growing, very little is known about its secondary structure within the whole grain, and previous research has primarily focused on oat isolates. It remains unclear whether there are differences in protein secondary structure among oat varieties, how rapidly it changes during steam pressure toasting treatment, and whether processing-induced molecular changes are accompanied by alterations in the tissue distribution of major macronutrients. To the best of our knowledge, no study has yet combined FTIR chemical imaging with synchrotron-based point spectroscopy to investigate varieties and steam-pressure toasting duration effects in oat grains. The objectives of this study were as follows: (i) to map the spatial distribution of proteins, carbohydrates, and lipids within the sections of oat seeds and to determine whether this distribution is a conserved characteristic of the grain or varies with oat varieties and SPT durations; (ii) to quantify the effects of oat varieties on the protein secondary structures; (iii) to determine the effects of SPT durations (0, 30, 60, 90, 120 min) on protein secondary structure and starch and lipid matrices; and (iv) to establish a transparent, reproducible workflow for chemical imaging, spectral preprocessing, and Amide I peak fitting analysis for synchrotron-based Mid-IR. We hypothesized that the influence of oat varieties on protein secondary structures was minor compared to SPT treatments, and SPT would drive the transition of proteins from disordered structures to ordered and aggregated conformations.

## 2. Materials and methods

### 2.1 Sample preparation

#### 2.1.1 Oat samples

Four cool-season oat varieties (CDC Haymaker, Summit, CDC Nasser, CDC Arborg) were provided by the Crop Development Center (CDC, Dr. Aaron Beattie) at the University of Saskatchewan, Canada. They were grown in the Crop Research Fields in Saskatchewan, Canada, and harvested at full maturity for commercial purposes over three consecutive years (2018, 2019, and 2020), serving as biological replicates (blocks) in a randomized complete block design (RCBD). In addition, a barley variety (CDC Austenson) obtained from the same source was used as a grain reference for spectral analysis. Two experiments were conducted in this study. The first experiment focused on the impact of oat variety on macronutrients (proteins, carbohydrates and lipids) distribution and protein secondary structures, using the four varieties listed above. The second experiment examined the impact of steam-pressure toasting (SPT) durations on macronutrients (proteins, carbohydrates and lipids) distribution and protein secondary structures of CDC Nasser oats.

#### 2.1.2 Steam-pressure toasting

For the processing experiment, CDC Nasser oat samples from three harvest years (2018, 2019, and 2020) were subjected to SPT treatment. Eight hundred grams of whole oat kernels were heated on an uncovered aluminum tray using a steam sterilizer (Amsco Century, Steris, Mentor, OH, USA), with an initial dry matter (DM) content of approximately 91.0%. The autoclave was operated in gravity circulation mode at 121□. Each complete cycle included the specified sterilization time (30, 60, 90 or 120 minutes), followed by 10 minutes of vacuum drying (10 inches of mercury), and 10 minutes of cooling. After removing the samples from the autoclave, the trays were left at room temperature, and the samples were then stored for subsequent analysis. Untreated samples served as the control (0 min). For the 2018 and 2020 harvest years, all five durations (0, 30, 60, 90 and 120 min) were prepared, whereas the 30 min duration was not available for the 2019 harvest year and four durations (0, 60, 90 and 120 min) were prepared. Steam-pressure toasting has been described as a method equivalent to autoclaving (Goelema, J. O., 1999; van del Poel et al., 2005), and these two terms have been used interchangeably in the literature (Aguilera et al., 1992; Goelema et al., 1998; Rodríguez Espinosa and Yu, 2025; Yu et al., 2000).

#### 2.1.3 Sample cryo-sectioning and mounting

For Transmission Imaging and synchrotron-based Mid-IR spectroscopy analysis, seed sections were prepared using the following cryo-sectioning protocol. Sections harvested from 2018 and 2019 were prepared in-house at the Earth Sciences Lab of the Canadian Light Source Inc., Saskatoon, Canada. For each variety (Experiment 1) and each SPT treatment (Experiment 2), these seeds were randomly selected from each harvest year and soaked overnight in tubes with ultrapure water to soften the tissue before sectioning (Deng et al., 2025, 2023b). After soaking for approximately 12 hours, the seeds were rapidly frozen using liquid nitrogen. Forceps were utilized to hold each sample in place, ensuring that the see was positioned as centrally as possible during freezing. Once the sample was completely frozen, the tube was held by hand for 1-2 minutes to allow it to thaw slightly. The frozen column containing the sample was then removed using forceps and attached to the sample holder of the metal cryostat using embedding medium (Leica Surgipath FSC 22 Clear Frozen Section Compound). Subsequently, a Leica CM1950 cryostat (Leica Microsystems, Wetzlar, Germany) was utilized to section the sample into 6 jim thick cross sections at a temperature of -20 . Sections from 2020 were prepared by an external sectioning service to the same nominal thickness, but without the overnight hydration step. All sections were unstained and mounted on BaF_2_ windows (1 mm thick, 13 mm in diameter; Crystran Ltd, Dorset, UK), which are transparent in the mid-infrared spectral range. A Leica S8 APO stereo-zoom microscope with a camera (Leica Microsystems, Germany) was used to collect microphotographs of the transverse sections. A representative section with high-quality cutting was selected from each kernel for subsequent analysis.

### 2.2 Transmission imaging

Fourier transform infrared (FTIR) transmission imaging was performed to map the spatial distribution of macronutrients (protein, carbohydrates, and lipids) within the oat seed cross-sections. Measurements were conducted at the Mid-IR beamline of the CLS, Saskatoon, Canada, using an Agilent Cary 670 FTIR spectrometer coupled with a Cary 620 FTIR microscope (Agilent Technologies, Santa Clara, CA, USA) [Figure S1 (a) and (b)]. The system was equipped with a globar (internal thermal) source for broadband illumination and a 128 × 128 element focal plane array (FPA) detector operating at cryogenic temperature (<80 K). A 15× objective was used for image collection, providing a pixel size of 5.5 jim. Transmission images were collected over the Mid-IR spectral range of 4000-800 cm^-1^ at 4 cm^-1^ resolution with 128 co-added scans. Data acquisition and instrument control were managed using Resolutions Pro Imaging Method Editor software (Agilent Technologies).

Before sample analysis, FPA detector was calibrated using the non-uniformity correction procedure to ensure uniform signal response is uniform across all detector elements. A clean area on the BaF^2^ window, free from particles or contamination, was then identified and used as the background reference position. Background spectra were collected at this position before each sample. A visible light image of the sample was captured to guide the selection of the infrared measurement area, which was defined by marking the region of interest encompassing the entire cross-section of the seed. Transmission infrared images were then collected as mosaic tiles covering the defined measurement area. The Agilent Cary 620 system offers a large field of view (up to 700 jim × 700 jim per tile) and features calibrated mosaic stitching capabilities, enabling effective large area chemical imaging of the entire seed cross-sections. The Supplemental Figure S2 presents the general workflow from sample preparation to data collection.

Three seeds were imaged for each oat variety from each harvest year (Experiment 1) and for each SPT treatment time (Experiment 2). Chemical distribution maps were generated and visualized by integrating the baseline absorption intensities in the following Mid-IR spectral regions using Quasar software: the lipid-related region (1770-1719 cm^-1^, including C=O ester stretching vibrations), the protein-related Amide I region (1700-1600 cm^-1^), and the carbohydrate-related region (1178-946 cm^-1^). To ensure comparability across all samples and treatments, standardized color-scale limits were applied to all chemical maps: 0-4 for lipid maps, 0-25 for protein maps, and 0-110 for carbohydrate maps (calculated in units of relative absorbance intensity). By color-coding and overlaying the protein (green), lipid (red) and carbohydrate (blue) channels, composite multichannel images were constructed to visualize the colocalization and relative abundance of macronutrients within seed tissues. Additional workflow details are presented in Supplemental Figure S3. Moreover, visible light microscopy images of seed cross-sections were also captured to display the complete seed morphology for anatomical analysis.

To validate the distribution of macronutrients revealed by the chemical map, spectra from target points were extracted from anatomically distinct tissue regions using FTIR transmission imaging data in Quasar software, including aleurone layer, crease region, embryo, endosperm, pericarp and sub-aleurone layer.

### 2.3 Protein secondary structures analysis using synchrotron based Mid-IR

#### 2.3.1 Spectral acquisition

To obtain high-resolution point spectra for protein secondary structure analysis, the sample sections were transferred to the Bruker Hyperion 3000 IR microscope equipped with a mercury cadmium telluride (MCT) single-element detector (Bruker Optics, Ettlingen, Germany), coupled with a synchrotron light source at the Mid-IR beamline of the CLS [Figure S1. (c) and (d)]. The high brightness and small effective source size of synchrotron radiation enable diffraction-limited spatial resolution, which is essential for collecting spectra from specific microstructural regions within the seed tissue.

Approximately thirty spot spectra were randomly selected from the aleurone layer and endosperm region of each seed cross-section. Sampling points were chosen based on an adequate infrared signal with well-defined Amide I and II bands to ensure high-quality spectra for band deconvolution. Spectra were collected in transmission mode through an aperture of 4.5 × 4.5 “m over the 4000-800 cm^-1^ range at 4 cm^-1^ resolution with 128 co-added scans. Background spectra were collected from a clean area on the BaF_2_ window under identical acquisition parameters.

#### 2.3.2 Spectra preprocessing and peak-fitting workflow

All spectral and peak-fitting analysis were performed using Quasar (1.13.1) software. Additional procedures using Quasar software are presented in Supplemental Figure S4. Data were organized by harvest year, with each year analyzed independently. For each year, the Concatenate function was used to combine the spectral files into a single table for all four oat varieties (Experiment 1, with 3 replicate seeds per variety, designated as A, B, and C) and all SPT treatment durations (Experiment 2, with 3 replicate seeds per treatment), with the data Source ID appended as a metadata attribute for sample identification. This process was repeated for three harvest years (2018, 2019, and 2020).

The concatenated spectra were first preprocessed through the following sequential steps: (1) the spectra were truncated to the fingerprint region (1800-900 cm^-1^); (2) Gaussian smoothing was applied with a standard deviation of 2; (3) baseline correction was performed using the rubber band method; (4) the spectra were normalized. Next, the preprocessed spectra for each sample were averaged using an average spectral function grouped by Source ID, with each seed producing a representative average spectrum (i.e., the average of approximately 30 spot spectra), resulting in 27 average spectra per year.

To verify the peak positions prior to deconvolution, parallel processing branches were applied to the average spectrum using a Savitzky-Golay filter (window size = 11, polynomial order = 2, derivative order = 2) to calculate the second-derivative spectrum. The second-derivative spectra were examined in the range of 1700-1500 cm^-1^ to confirm the positions of the component spectral bands and to identify any spectra exhibiting anomalous features. Anomalous spectra or those deviating significantly from most of the spectra were selected for separate peak-fitting.

#### 2.3.3 Amide I band deconvolution

Peak-fitting deconvolution was performed on the averaged spectra truncated to the 1700-1500 cm^-1^ region, containing the Amide I (1700-1600 cm^-1^) and Amide II (1600-1500 cm^-1^) bands. A total of nine Gaussian subpeaks were fitted, including 6 in the Amide I region and 3 in the Amide II region. The center, amplitude, and o (width) parameters of each Gaussian peak were adjusted until the sum of all fitted subpeaks closely matched the original experiment spectrum. The detailed parameters were listed in Table 1.

Based on established spectral assignments, the six Amide I subpeaks were assigned to protein secondary structure components according to their center wavenumber positions (Byler and Susi, 1986; Dong et al., 1990; Ma et al., 2001): 0-sheet structures (1640-1600 cm^-1^), represented by sub-peaks at approximately 1614 and 1636 cm^-1^; random coil (1650-1640 cm^-1^), represented by a sub-peak at approximately 1645 cm^-1^; a-helix (1660-1650 cm^-1^), represented by a sub-peak at approximately 165 3 cm^-1^; and β-turn structures (1700-1660 cm^-1^), represented by sub-peaks at approximately 1661 and 1684 cm^-1^. The relative proportion of each secondary structure was calculated as the sum of the integrated areas of its corresponding subpeaks divided by the total sum of the integrated areas of all six Amide I subpeaks, expressed as a percentage.

#### 2.3.4 Sub-peak curve reconstruction for visualization

In the Quasar software, the peak-fitting outputs provide fitting parameters (center, amplitude, sigma, and area) for each Gaussian subpeak and the overall fitted curve. However, it does not directly export individual subpeak curves as functions of wavenumber. To generate the individual subpeak spectra required for visualization in Origin software (OriginLab Corporation, Northampton, MA, USA), a scaling decomposition method was employed. For each sample, the Gaussian function was utilized to reconstruct the raw Gaussian curve for each subpeak *i* from its fitting parameters:

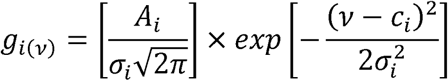

where *A_i_* is the integrated area, *c*_i_ is the center wavenumber, and w is the sigma (standard deviation) of subpeak *i*, and v is the wavenumber. The total reconstructed sum of all subpeaks was then calculated as:

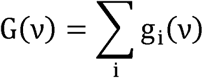

Since the absolute intensities of the reconstruction and G(v)do not always match the actual fitted curve F(v) derived from Quasar software (due to differences in internal scaling and normalization within the software), a proportional decomposition method is employed. The fractional contribution of each subpeak at each wavenumber was calculated as:

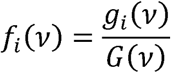

The scaled subpeak curve was then obtained by multiplying this fraction by the actual fitted total from Quasar:

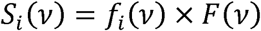

This method ensures that the sum of all scaled subpeaks is exactly equal to the fitted curve at every wavenumber point [i.e., *Z* S_i_ (v)= F (v)], while preserving the relative shape and proportions of each component. The residual between the raw experiment spectrum and the fitted curve was calculated as:

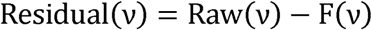

For each sample, the output data table included wavenumbers, raw spectra, the total number of fits, individually scaled subpeak curves, and residuals. Plotting in Origin software was performed using the output data table.

### 2.4 Data analysis

Both experiments employed a randomized complete block design (RCBD), with harvest year (2018, 2019, and 2020) treated as blocks to account for annual environmental variations. For each treatment in each year, three seed sections (subsamples) were measured, and their average was used as the block-level observation, yielding three observations (n=3 blocks) per treatment. The model for the RCBD is expressed as:

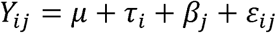

where *Y_ij_* is the observed response variable for the *i*th treatment in the *j*th block, µ is the overall population mean, τ_i_ is the fixed effect of the treatment (*i* = 1, 2,…, *a*; where *a* = 4 in Experiment 1 and *a* = 5 in Experiment 2),*β*_j_ is the random effect of the *j*th block (*/’* = 1, 2, 3, corresponding to the harvest years 2018, 2019 and 2020), and ε_ij_ is the random error term, assumed to be independently and normally distributed with mean zero and constant variance. In Experiment 1, the treatment factor was oat variety across four levels, including CDC Haymaker, Summit, CDC Nasser, and CDC Arborg. In Experiment 2, the treatment factors were 0 (untreated control), 30, 60, 90 and 120 minutes of SPT. The response variables for both experiments were the relative proportions (%) of the four protein secondary structure components, including α-helix, β-sheet, random coil, β-turn, and the ratio of α-helix to β-sheet.

Analysis of variance (ANOVA) was utilized to test the significant treatment effects. The null hypothesis (H*ø*:τ_1_=τ_2_=…=τ_a_=0) against the alternative hypothesis (H*1*: at least one . ≠ 0) was tested at a significance level of a = 0.05. When the overall F-test indicated a significant treatment effect (*P* < 0.05), Fisher’s protected LSD test was used for pairwise comparisons to identify specific treatment differences. Results of multiple comparisons were presented using letters, where treatments sharing the same letter showed no significant differences at *P* < 0.05.

All statistical analysis were performed using Python 3.12 and the statsmodels package (version 0.14) to conduct analysis of variance (ANOVA) and LSD’s test. The matplotlib package (version 3.9) was used to generate publication-quality figures, including bar charts with standard error of the mean (SEM) bars, individual data points representing the variability of the three block-level observations, letter annotations, and overall model P-value annotations. Data were exported in PNG and TIFF formats at a resolution of 600 DPI.

## 3. Results and discussion

### 3.1 Impact of oat variety on macronutrient distribution and protein secondary structure 3.1.1 Spatial distribution and spectral characterization of macronutrients

Figure 1 and Figure 3 reveal the spatial distribution of macronutrients within the seed cross-sections using FTIR Transmission Imaging from the 2018 and 2020 harvesting years, respectively. Across all four oat varieties, carbohydrates were predominantly localized in the endosperm region, presenting the highest absorbance intensity (up to ∼110 absorbance units). Proteins were mainly concentrated in the aleurone layer, sub-aleurone layer and embryo regions, with relatively lower concentrations detected within the endosperm matrix. Additionally, lipids were primarily found in the crease region and embryo with contributions from the aleurone layer and the outer pericarp, and minor concentrations were also observed in the endosperm. Notably, the spatial distribution of these macronutrients was highly consistent, with no apparent visual changes in the distribution patterns across CDC Haymaker, Summit, CDC Nasser, and CDC Arborg. The same overall pattern was also similarly in the images of the two additional replicate samples provided in the supplemental materials (Figure S5 and S6). These spatial distributions were consistent with previous synchrotron-based infrared studies of cereal and legume seeds, in which carbohydrates, proteins and lipids were similarly distributed in the endosperm, aleurone layer and embryo, as well as crease and pericarp regions, respectively (Feng et al., 2020; Yu et al., 2004b). These observations suggest that the distribution of macronutrients is a conservative structural trait of oat seeds rather than a variety dependent characteristic. However, small differences in intensity exist among varieties, which may reflect genotype related differences in nutrient accumulation density. Since the absorbance in transmission imaging also depends on the section thickness, these differences in intensity should be interpreted with caution. They were not used here as a quantitative measure of nutrient concentration.

**Figure 1.**
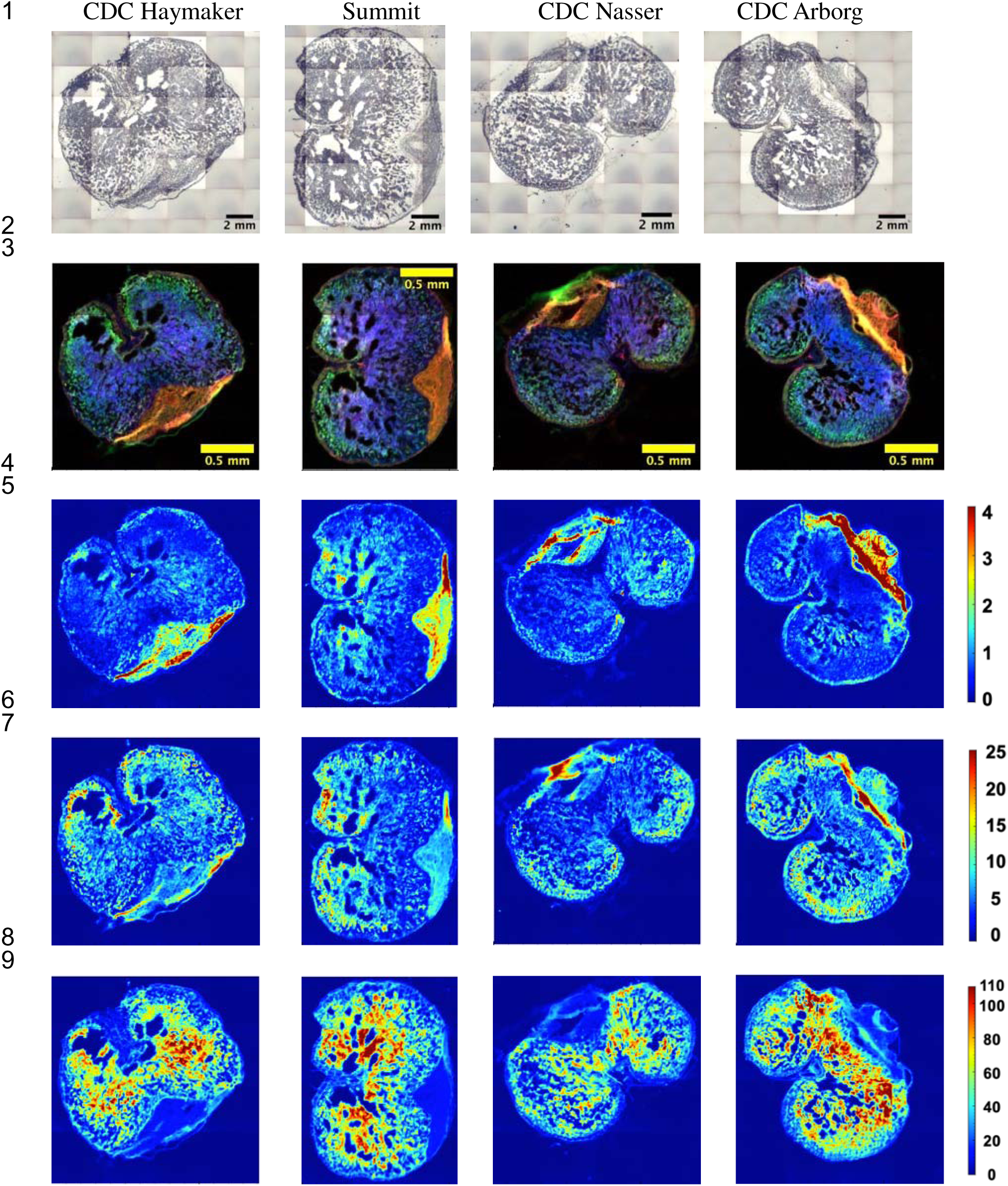
Spatial distribution of macronutrients (lipids, protein, carbohydrates) within the seed cross-sections of four cool-season oat varieties (CDC Haymaker, Summit, CDC Nasser, CDC Arborg) harvested from Year 2018. **Row 1:** Visible light microscopy images displaying the intact seed morphology. **Row 2:** Composite multi-channel images illustrating the combined distribution of protein (green), carbohydrate (blue) and lipid (red). **Rows 3-5:** Individual chemical heat maps for the lipids, protein and carbohydrates distributions, respectively. The color scales represent the relative absorbance intensity, where warmer colors denote higher concentrations.

**Figure 2.**
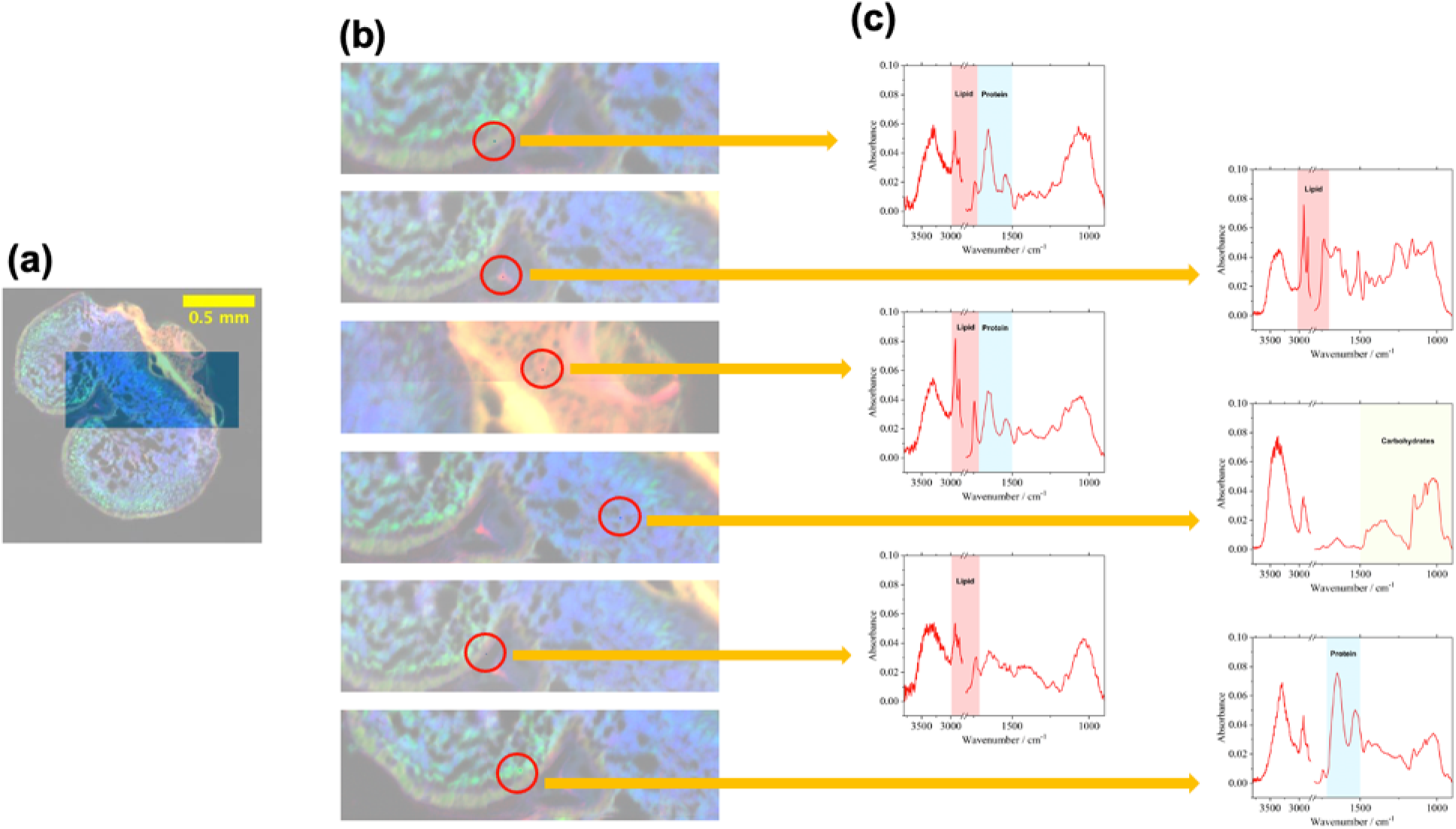
Spatial and spectral analysis of CDC Arborg harvested from Year 2018. **(a)** Mid-IR image of the intact seed cross-section, with shaded rectangular area highlighting the target**ed** region of interest for sampling. **(b)** Magnified view of the selected region. Red circles indic**ate** specific spot samples located on the major anatomical structures: aleurone layer, crease region, embryo, endosperm, pericarp, sub-aleurone layer. **(c)** Extracted Mid-IR absorbance spectra **f**or each spot, illustrating the localized chemical profiles and characteristic bands of carbohydra**te,** lipid, and protein.

**Figure 3.**
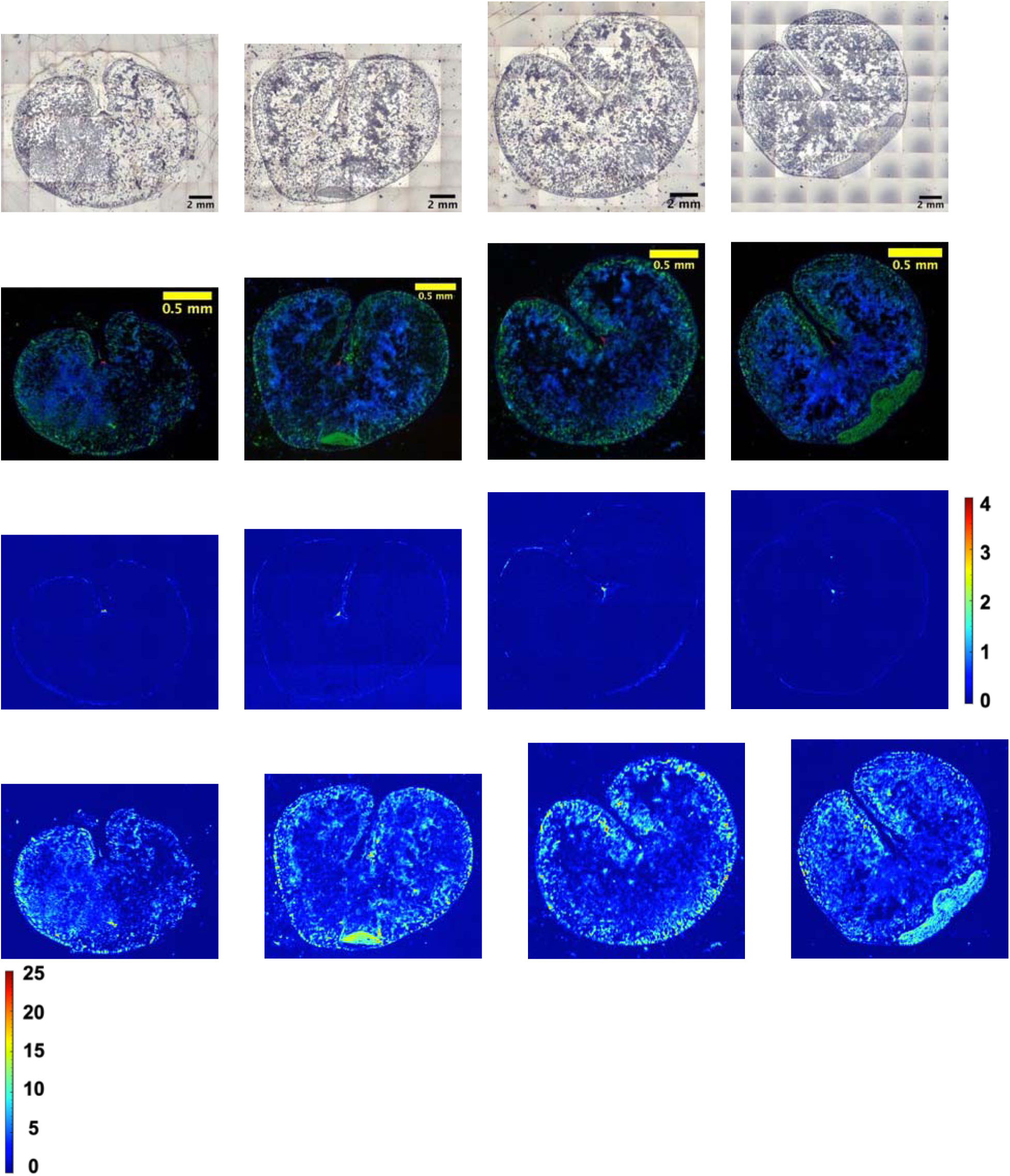
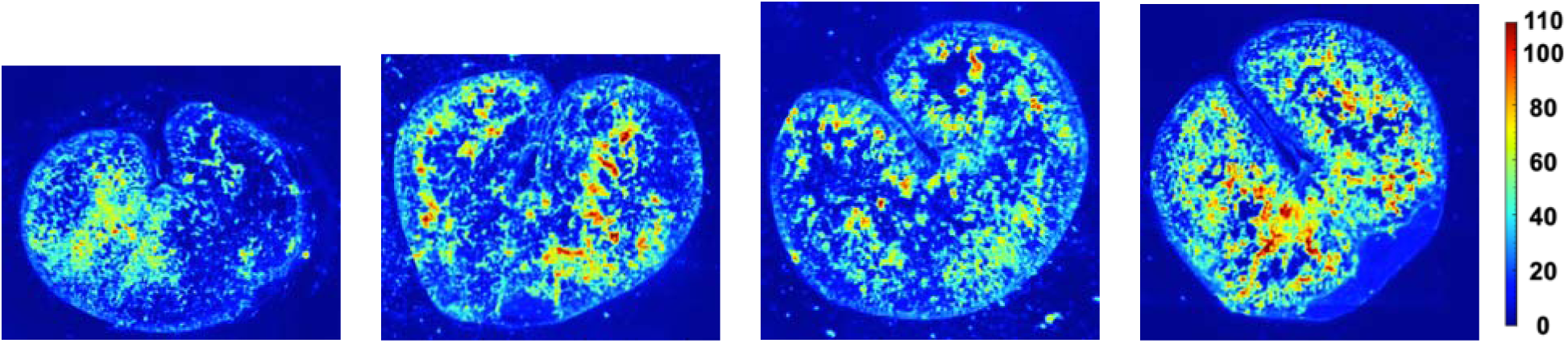
Spatial distribution of macronutrients (lipids, protein, carbohydrates) within the see**d** cross-sections of four cool-season oat varieties (CDC Haymaker, Summit, CDC Nasser, CD**C** Arborg) harvested from Year 2020. **Row 1:** Visible light microscopy images displaying the int**ac**t seed morphology. **Row 2:** Composite multi-channel images illustrating the combined distribution of protein (green), carbohydrate (blue) and lipid (red). **Rows 3-5:** Individual chemical heat ma**ps** for the lipids, protein and carbohydrates distributions, respectively. The color scales represent t**he** relative absorbance intensity, where warmer colors denote higher concentrations.

There is a significant difference in the overall signal intensity of the chemical maps from 2018 (Fig. 1) and 2020 (Fig. 3). The samples from 2018 were soaked overnight in ultrapure water prior to cryo-sectioning (see Section 2.1.3) and presented distinct spatial boundaries and strong absorption signal in all three maps of macronutrients. However, samples from 2020 were prepared by the external sectioning service without an overnight hydration step and showed markedly reduced signal intensity. In particular, the spaces between tissues were less clearly defined in the lipid and protein images. This difference may be due to the lack of tissue hydration prior to sectioning. Without sufficient softening, these dry and brittle seed tissues were more susceptible to mechanical damage, fragmentation, and uneven section thickness during cryo-sectioning, all of which may affect the nutrient distribution patterns. The samples from 2019 were also prepared using the same overnight hydration protocol, and their distribution patterns were similar to those of the samples from 2018 (Supplemental Fig. S8), demonstrating that overnight soaking step helps to preserve the natural spatial organization of macronutrients during section preparation. Therefore, the chemical maps of the hydrated samples (2018) served as the primary reference for spatial distribution analysis during this study, while the maps of samples from 2020 exhibited the consistency of the basic distribution patterns from year to year, despite differences in sample preparation.

To validate the spatial distribution patterns identified through chemical imaging, targeted spot-sampling was conducted on CDC Arborg harvested in 2018 across six different anatomical regions spanning the aleurone layer, crease region, embryo, endosperm, pericarp, and sub-aleurone layer (Figure 2). The corresponding visible light microscopy images confirmed the morphological characteristics of each tissue sample. The Mid-IR absorbance spectra obtained from each spot exhibited characteristic spectral curves consistent with their expected biochemical compositions and with the established band assignments for proteins, lipids and carbohydrates (Barth, 2007; Liu and Yu, 2016; Wetzel et al., 1998). The endosperm spots displayed strong and distinct absorption bands in the carbohydrate region (1500-900 cm^-1^), dominated by C-H bending, C-C stretching, P=O stretching, and C-O/C-O-C stretching modes, with minimal contributions from protein or lipid bands. The spectra of the crease region and pericarp showed distinct lipid-related features, including CH_2_ and CH_3_ stretching (∼3000-2800 cm^-1^) and C=O stretching (∼1740 cm^-1^). The spectra of the aleurone layer and sub-aleurone layers showed distinct Amide I (∼1650 cm^-1^) and Amide II (∼1550 cm^-1^) absorption bands, confirming the protein-rich nature of these tissues. Additionally, the embryo spectra also exhibited distinct protein and lipid absorption bands. These tissue-specific spectral band assignments reproduced the reported evaluation of chickpea seed tissues at a high spatial resolution using synchrotron infrared spectroscopy (Feng et al., 2020). Compared to spot-sampling for oats harvested in 2020 (Figure 4), the results showed consistent tissue-specific spectral profiles; hence, the endosperm was dominated by the carbohydrate band, the crease region dominated by the lipid band, the aleurone layer dominated by the protein band, and the embryo dominated by the protein and lipid bands, although the overall signal intensity decreased. These results confirm that the fundamental biochemical characteristics of each tissue type are preserved regardless of the sample preparation method. In conclusion, these spectral results support the chemical imaging findings and provide point-level spectral evidence for the tissue-specific distribution of macronutrients in oat seeds.

**Figure 4.**
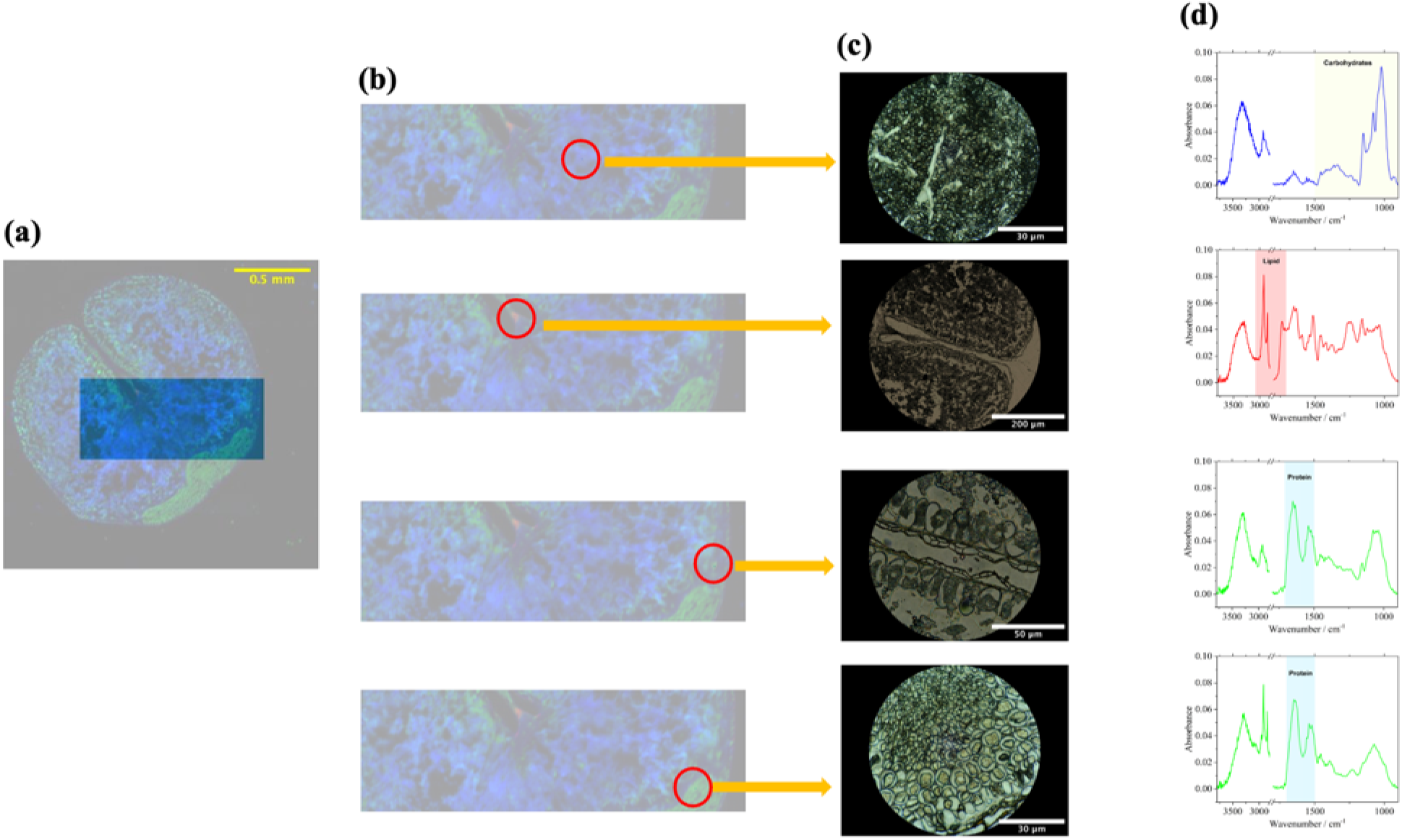
Spatial and spectral analysis of CDC Arborg harvested from Year 2020. **(a)** Mid-IR image of the intact seed cross-section, with shaded rectangular area highlighting the target**ed** region of interest for sampling. **(b)** Magnified view of the selected region. Red circles indic**ate** specific spot samples located on main anatomical structures: endosperm, crease region, sub**­**aleurone layer and embryo. **(c)** Visible light microscopic images corresponding to each of th**e** selected sampling spots. **(d)** Extracted Mid-IR absorbance spectra for each spot, illustrating th**e** localized chemical profiles and characteristic bands of carbohydrate, lipid, and protein.

#### 3.1.1 Global Mid-IR spectral profiles of four oat varieties and barley

The global synchrotron-based Mid-IR absorbance spectra (3600-900cm^-1^) of four oat varieties and one barley variety (CDC Austenson) presented overall similar spectral profiles, indicating a broadly biochemical composition across these cereal grains (Figure 5). All spectra showed three high-resolution spectral regions corresponding to the major lipid-related region (∼3000-2800 cm^- 1^ and ∼1740 cm^-1^), the protein-related region (∼1700-1500 cm^-1^) and the carbohydrate-related region (1500-900 cm^-1^). In the carbohydrate region, four oat varieties and barley variety displayed largely overlapping absorption profiles. The most notable difference was observed in the peak at approximately 1000 cm^-1^; therefore, peaks for the CDC Nasser, CDC Arborg, CDC Austenson barley were relatively smooth, while those for Summit and CDC Haymaker were sharper and more jagged. Because the shape of this region reflects the balance between ordered and amorphous starch domains (Sevenou et al., 2002; Van Soest et al., 1995), these differences may indicate subtle variations in the short-range order of starch among genotypes. In the protein region, subtle differences were observed in the relative intensities of the Amide I and Amide II bands among varieties, with CDC Austenson barley and CDC Arborg showing marginally higher protein-related absorbance. In the lipid region, the CH_2_/CH_3_ stretching and C=O stretching bands exhibited minor intensity variations among oat varieties. CDC Austenson barley displayed less pronounced lipid features (the C=O peak at 1740 cm^-1^), which was consistent with the generally lower oil content of barley compared to oats. These variety-dependent spectral differences are of practical relevance, as synchrotron-based FTIR characterization of barley endosperm has previously been linked to its rumen degradation characteristics (Yu et al., 2004a), suggesting that the subtle spectral variations observed among varieties in this study may likewise reflect differences in their nutritional and feed value.

**Figure 5.**
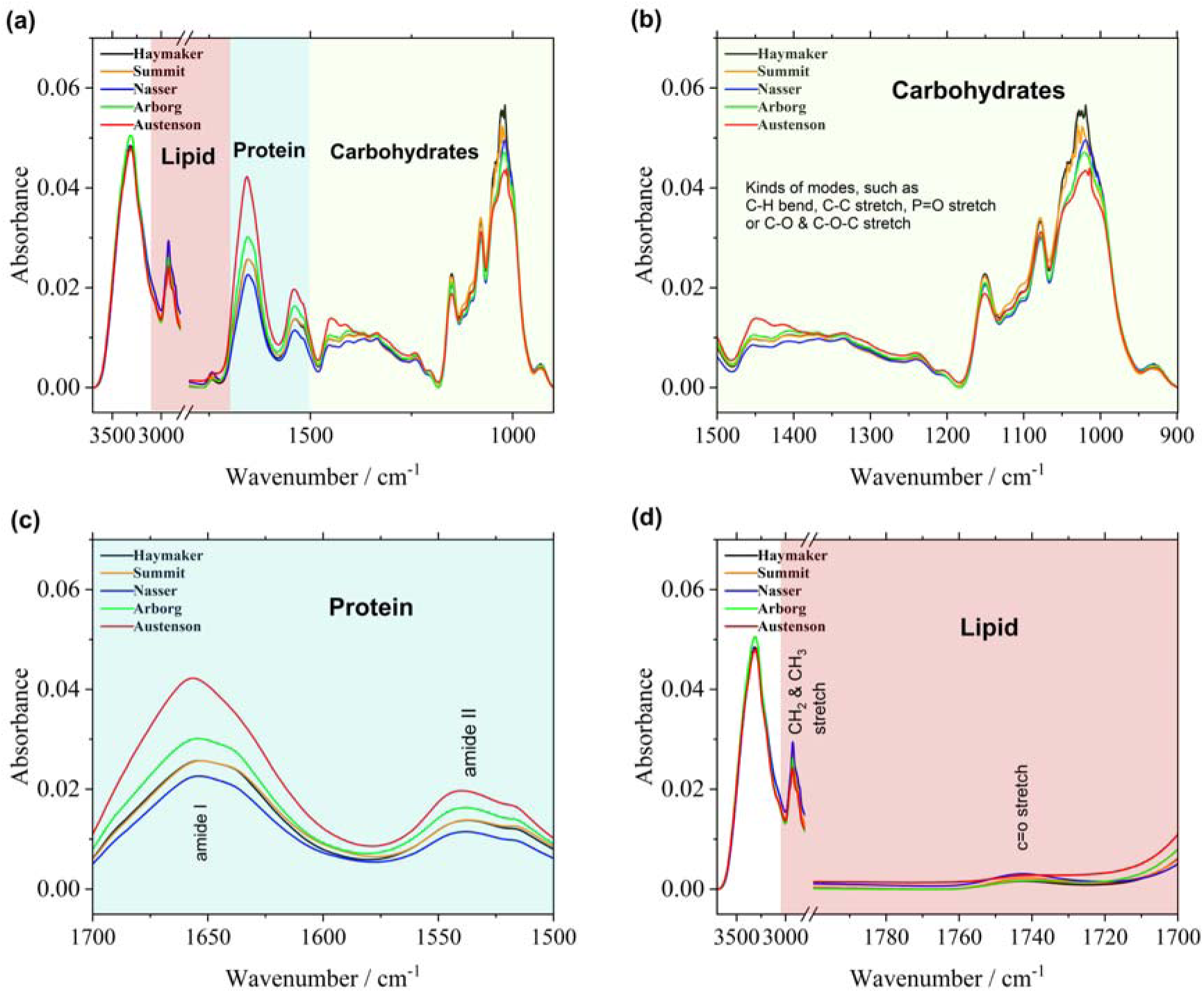
Global Mid-IR absorbance spectra of four cool-season oat varieties (CDC Haymaker, Summit, CDC Nasser, and CDC Arborg) and one barley variety (CDC Austenson). **(a)** The full raw Mid-IR spectral profile (approx. 3600-800 cm^-1^). The spectra for the oat varieties represent the weighted average of three biological replicates and then the average of three consecutive harvest years (2018, 2019 and 2020), while the barley spectrum represents the average of three replicates as reference. **(b)** Magnified view of the carbohydrate-associated region (1500-900 cm^- 1^), highlighting absorption modes such as C-H bending, C-C stretching, P=O stretching, and C- O/C-O-C stretching. **(c)** Magnified view of the protein-associated region (1700-1500 cm^-1^), detailing the distinct Amide I and Amide II absorption bands. **(d)** Magnified view of the lipid- associated region, displaying characteristic peaks for CH2 and CH3 stretching (∼3000-2800 cm^-1^) and C=O stretching (∼1740 cm^-1^) to facilitate the comparison of spectral differences.

#### 3.1.2 Protein secondary structure composition among oat varieties

To clarify the specific variations within the protein matrix, the Amide I band (1700-1600 cm^-1^) was deconvoluted to resolve potential secondary structural conformations among four oat varieties harvested in 2018. Barley grain was used as the reference (Figure 6). The fitting plots for the four oat varieties harvested in 2019 and 2020 are included in Supplemental Figure S10. The fitted curves matched the raw data closely. Curve-fitting resolved the overlapping contributions of individual protein secondary structure elements, with a-helix (pink), 0-sheet (purple and cyan), *β*-turn (orange and yellow), and random coil (green) clearly identified as well. The Gaussian band shapes were utilized throughout the study after comparing the Gaussian and Lorentzian multi-component models for the analysis of Amide I plant seed as reported by Yu (2005). Across all the oat varieties and barley, the overall Amide I band was mainly contributed by *β*-sheet and *β*-turn, reflecting the typical secondary structure composition of cereal proteins. This was similar to the findings reported by Liu et al. (2009), who identified approximately 74% *β*-sheets, 19% a-helices, and 7% *β*-turns in oat protein fractions using Amide I deconvolution. Analysis of subpeaks reveals differences in the relative contributions of these secondary structural components among diverse varieties, providing a basis for subsequent quantitative comparisons.

**Figure 6.**
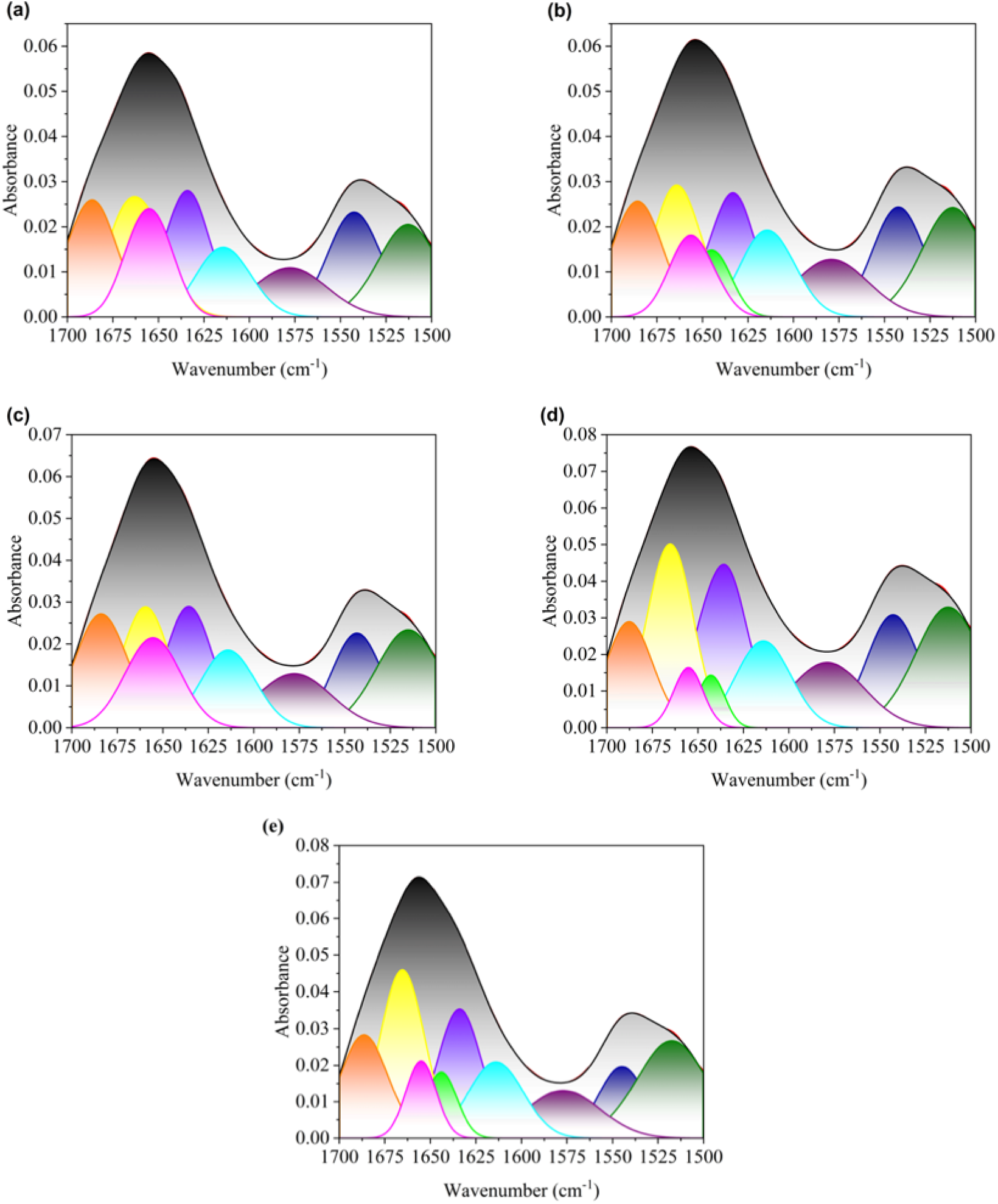
Peakfitting deconvolution of the Mid-IR Amide I region (1700-1600 cm-1) for determining the protein secondary structures of four cool-season oat varieties harvested in Year 2018 and one barley. The panels stand for (a) CDC Haymaker, (b) Summit, (c) CDC Nasser, (d) CDC Arborg and (e) CDC Austenson barley. In each panel, the black solid line represents the original raw spectrum calculated based on three replicates, and the red solid line indicates the overall fitted curve. The color-filled subpeaks under the curve correspond to protein secondary structures based on specific wavelength ranges: β-turn structures (1700-1660 cm-1) are represented by the orange (∼1684 cm^-1^) and yellow (∼1661 cm^-1^) peaks; the α-helix structure (1660^-1^650 cm^-1^) is represented by the pink (∼1653 cm^-1^) peak; the random coil (1650^-1^640 cm^-1^) is represented by the green (∼1645 cm^-1^) peak; and β-sheet structures (1640^-1^600 cm^-1^) are represented by the purple (∼1636 cm^-1^) and cyan (∼1614 cm^-1^) peaks.

Figure 7 shows relative proportions of protein secondary structures in the four cool-season oat varieties quantified by peak-fitting deconvolution in Amide I band. ANOVA analysis revealed statistically significant differences in a-helix proportion (*P = 0.026*), 0-turn proportion (*P = 0.047*), and a-helix/0-sheet ratio (*P = 0.048*) among four oat varieties. In contrast, no significant differences were detected for *β*-sheet (*P = 0.769*) and random coil (*P = 0.377*) proportions. Multiple comparison test (LSD) identified specific pairwise differences among the four oat varieties. CDC Nasser and Summit had significantly higher a-helix proportions than CDC Haymaker and Arborg (*P < 0.05*). For *β*-turn proportion, CDC Haymaker was significantly higher than that in Summit, while CDC Nasser and Arborg were intermediate. The ratio of a - helix to *β*-sheet was similar to that of a-helices, but the ratio in CDC Nasser was significantly higher than that in CDC Arborg, indicating a greater relative contribution of a-helix structures in CDC Nasser oats. These findings suggest that the protein secondary structure composition is broadly consistent among four oat varieties, but genotype-related variations exist, particularly in the ratios of *a* -helix to *β*-sheet. CDC Nasser and Summit tend to have higher *a* -helix content, which may be related to protein functionality and nutritional quality. In feed research, the ratio of α-helix to *β*-sheet was used as an indicator of protein accessibility to digestive enzymes (Carbonaro et al., 2012; Yu et al., 2004b).

**Figure 7.**
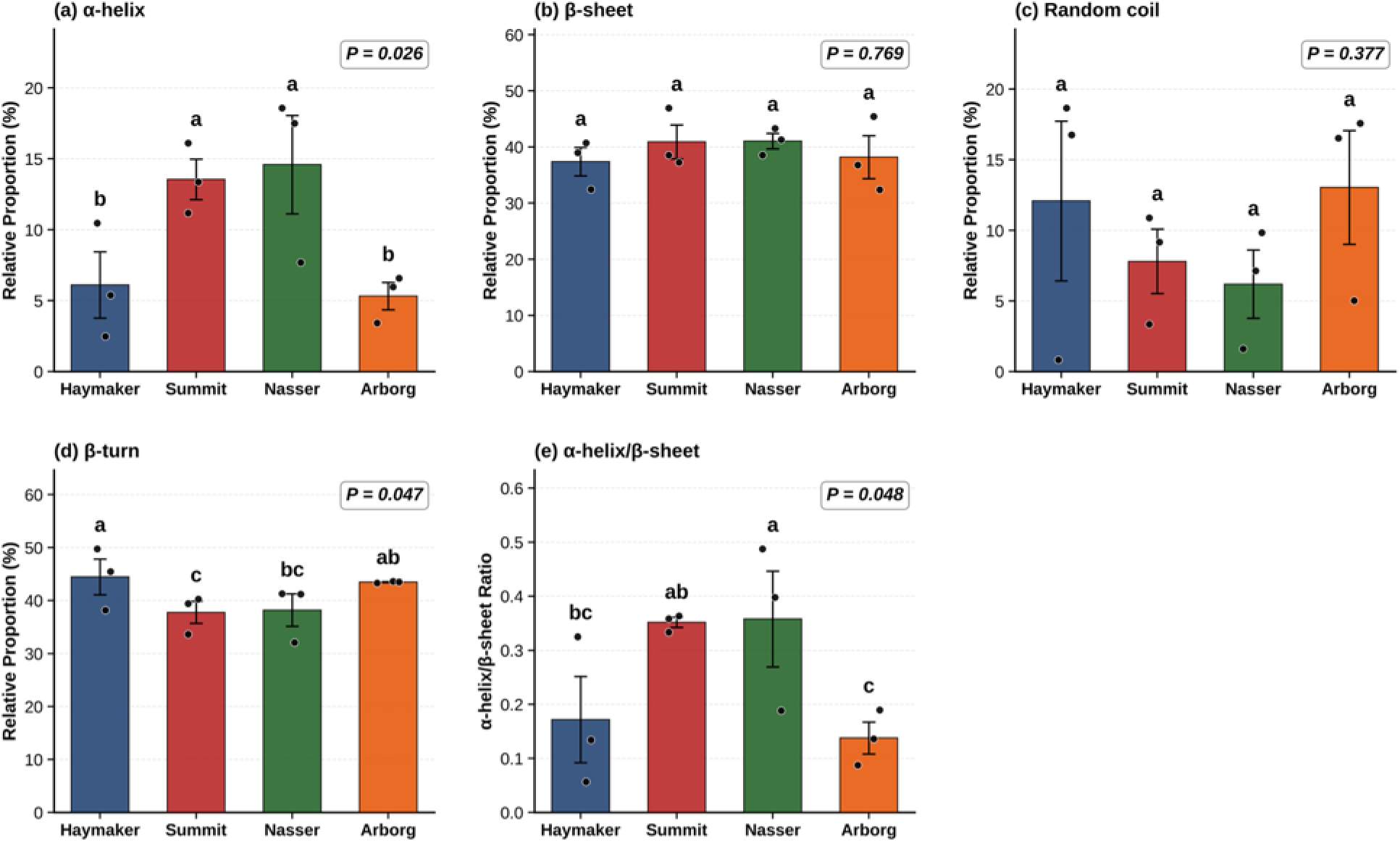
Relative proportions of secondary protein strutures among four cool-season o**a**t varieties (CDC Haymaker, Summit, CDC Nasser, and CDC Arborg). The quantitative resu**l**ts were derived from the peak-fitting deconvolution of the synchrotron-based Mid-IR Amide I ban**d**. The panels present the calculated percentage for **(a)** -helix, **(b)** -sheet, **(c)** random coil, **(d) -** turn, **(e)** the ratio of -helix to -sheet. For each variety within each harvest year, three se**e**d cross-sections (sub-samples) were measured and averaged to obtain a single block-lev**e**l observation. The bar heights represent the mean value of three harvesting years, and t**he** individual black dots indicate the block-level means from 2018, 2019 and 2020 (n = 3). Er**r**or bars represent the standard error means (SEM). ANOVA analysis was performed under **a** randomized complete block design (RCBD) with harvesting year as the blocking factor. Over**a**ll model *P*-values are displayed in the upper right corner of each panel. Means with the differe**n**t letters are significantly different (*P < 0.05*), and multi-treatment comparison is LSD method.

It is worth noting that the *β*-sheets observed in this study, which dominated the Amide I spectrum, differ from the findings of an earlier synchrotron-based infrared study by Yu et al. (2004b). In that report, oat endosperm tissue was found to contain approximately 92% a-helix and 2% *β*-sheet. Upon closer comparison, the difference appeared to be methodological rather than biological. First, Yu et al., (2004b) reported only two components (α-helix and 0-sheet) and expressed them as percentage of these two components alone, whereas the present work fitted six

Amide I subpeaks and normalized each conformation to the total area of Amide I, therefore *β* - turns and random coils were also part of Amide I. Second, the band-fitting functions were different: they used the Lorentz function, while the Gaussian function was utilized throughout this study, following the report that Gaussian modeling fitted the multi-peak curves of protein secondary structures more accurately than Lorentzian modelling (Yu, 2005). Furthermore, the number of subpeaks fitted, the constraints placed on their centers and widths, and the smoothing and baseline processing applied prior to fitting, all these factors may influence the resulting ratios, as Amide I decomposition might be particularly sensitive to these factors (Barth, 2007; Surewicz et al., 1993; Yu, 2005). The results of this study were more consistent with those reported by Liu et al. (2009) for oat protein isolate. Therefore, the absolute proportions reported here should not be interpreted as constants at the species level. Comparisons among varieties and treatments must be obtained using the same fitting procedure to provide a more reliable basis for interpretation.

### 3.2 Impact of steam-pressure toasting (SPT) on macronutrient distribution and protein secondary structure

#### 3.2.1 Effect of SPT durations on macronutrient distribution and Mid-IR spectral features

The spatial distribution of macronutrients in CDC Nasser oat seed cross-sections at five steam­pressure toasting (SPT) durations (0, 30, 60, 90 and 120 min) was shown in Figure 8 (harvested in 2018) and Figure 10 (harvested in 2020), with replicate images provided in the supplemental materials (Figure S7 and S9, respectively). Additionally, the spatial distribution of macronutrients in CDC Nasser oat seeds from the year harvested in 2019 at four SPT durations (0, 60, 90 and 120 min) were presented in Supplemental Figure S8, including three replicates. Across all processing durations, the basic spatial distribution of macronutrients remained unchanged. Lipids were mainly localized in the crease region and the embryo, with minor contributions from the aleurone layer and pericarp. Protein was primarily concentrated in the aleurone layer, sub-aleurone layer and embryo, with lower levels found in the endosperm. Carbohydrates were primarily distributed in the endosperm. These distribution patterns were consistent with the results observed among varieties (Fig. 1), confirming that the tissue-specific localization of macronutrients remained structurally robust and resistant to redistribution under SPT treatment conditions.

**Figure 8.**
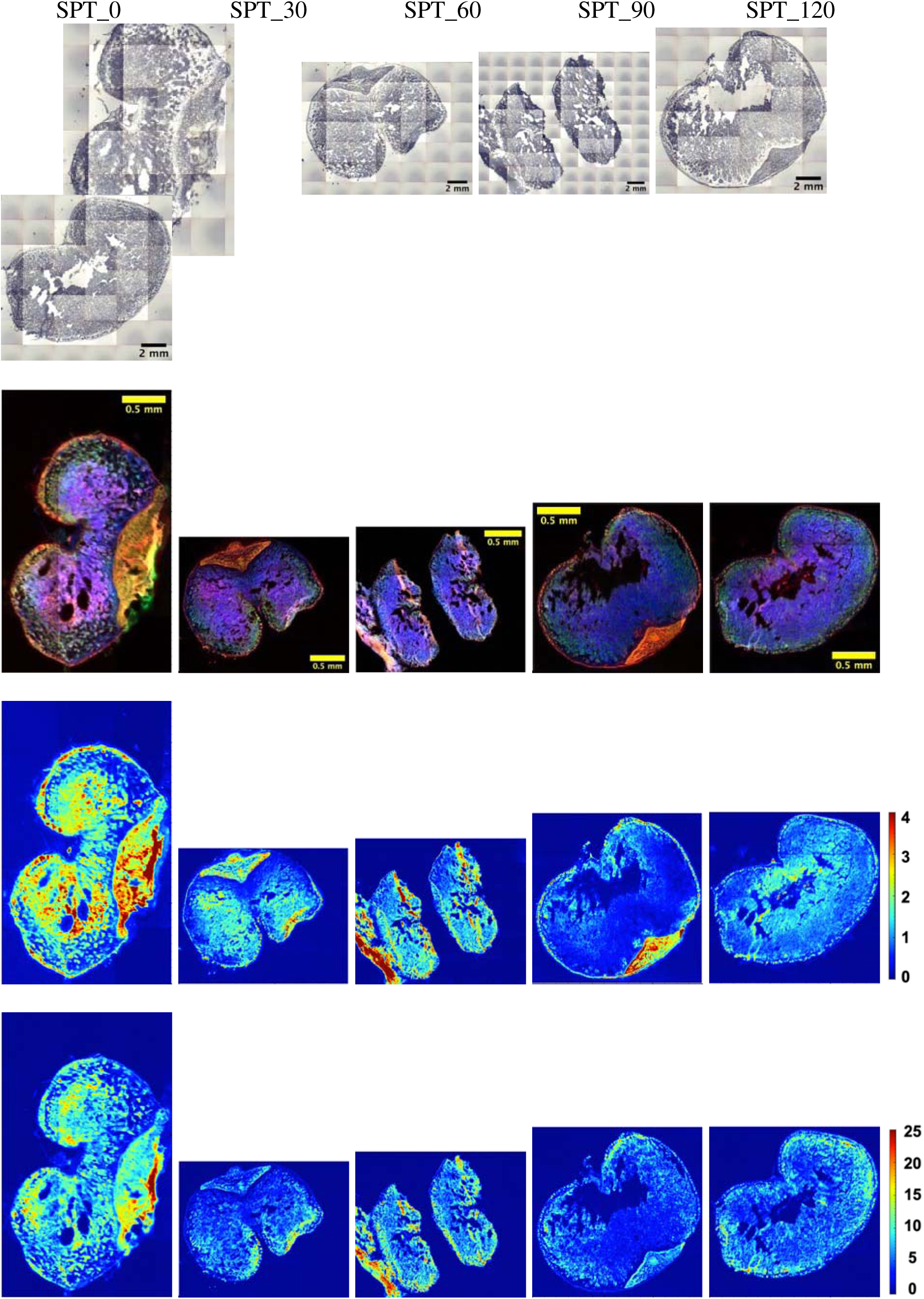
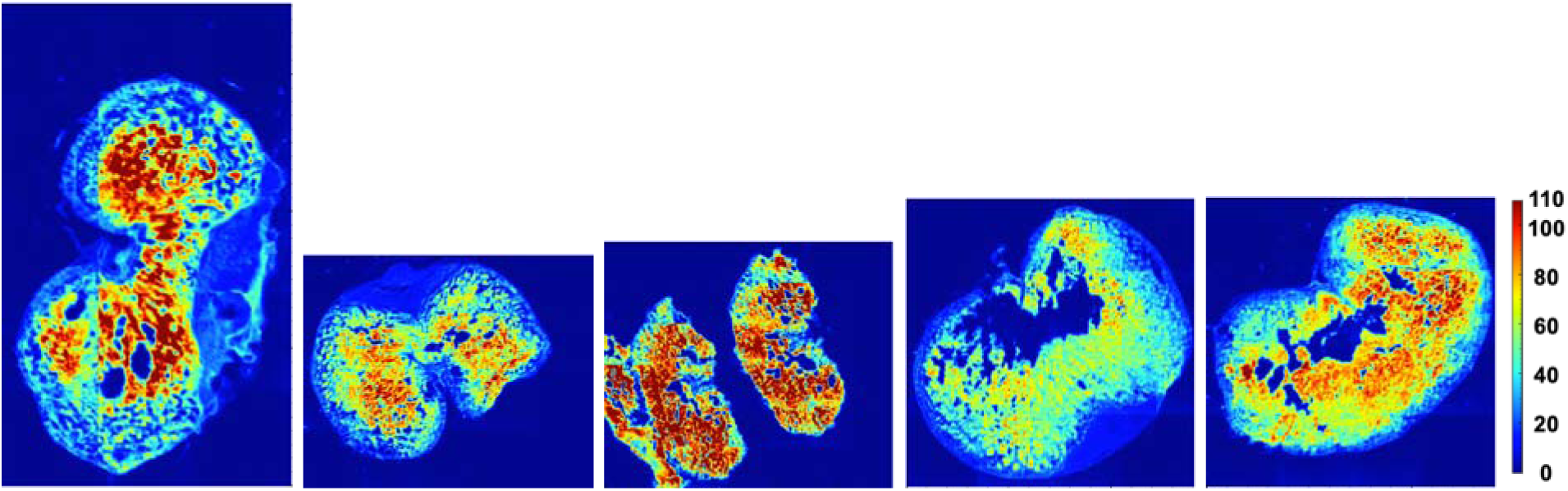
Spatial distribution of macronutrients within the seed cross-sections of the CD**C** Nasser oat with different durations of steam-pressre toasting (0, 30, 60, 90, and 120 minut**e**s) harvested from Year 2018. **Row 1:** Visible light microscopy images showing the intact see**d** morphology across the treatment timeline. **Row 2:** Composite multi-channel images depictin**g** the integrated distribution of proteins (green), lipids (red), and carbohydrates (blue). **Row 3-5:** Chemical heat maps illustrating the individual spatial distribution of lipid, protein, and carbohydrate, respectively. The color scales represent the relative absorbance intensity, whe**re** warmer colors denote higher concentrations. SPT: steam-pressure toasting.

**Figure 9.**
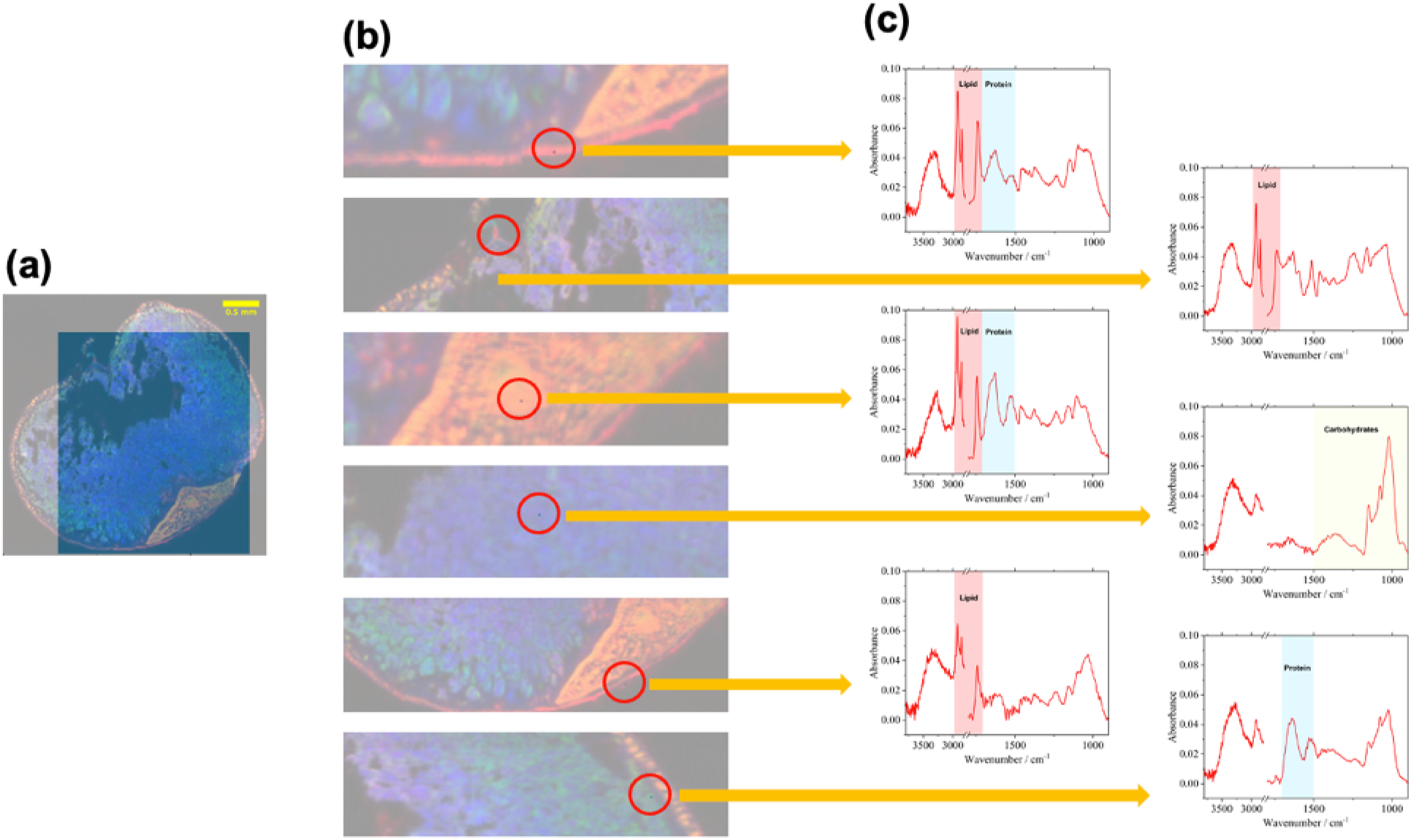
The spatial and spectral analysis of a CDC Nasser oat seed with 90 minutes of steam**­**pressure toasting harvested from Year 2018. **(a)** Mid-IR image of the intact seed cross-sectio**n,** highlighting the region of interest for targeted spot sampling. **(b)** Magnified view of the select**ed** region, with red circles indicating specific sampling spots in the main anatomical structure**s:** aleurone layer, crease region, embryo, endosperm, pericarp, and sub-aleurone layer. **(c)** Extracte**d** Mid-IR absorbance spectra for each spot, illustrating the localized chemical profiles **of** carbohydrates, lipids and proteins.

**Figure 10.**
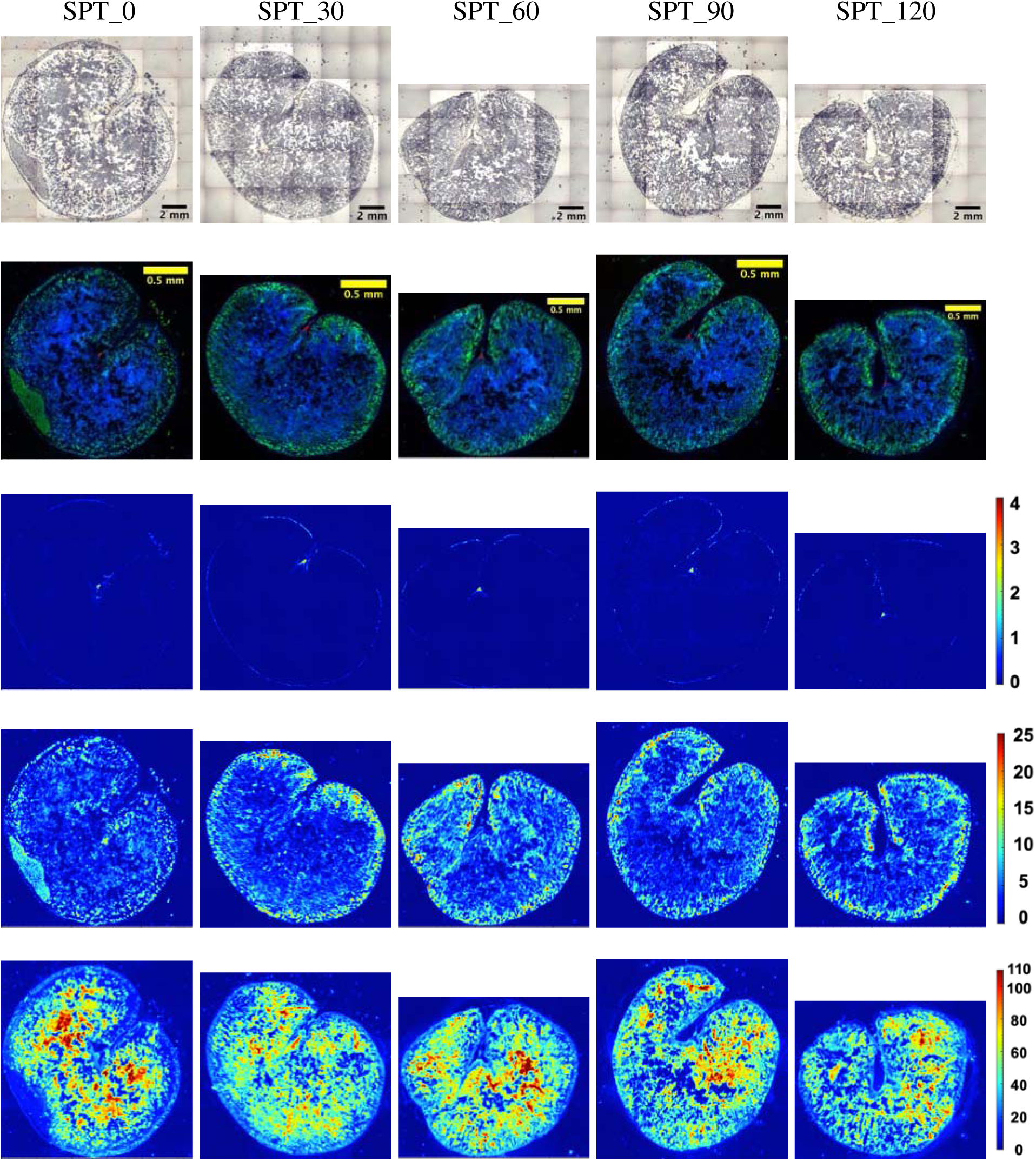
Spatial distribution of macronutrients within the seed cross-sections of the CDC Nasser oat with different durations of steam-pressure toasting (0, 30, 60, 90, and 120 minut**e**s) harvested from Year 2020. **Row 1:** Visible light microscopy images showing the intact see**d** morphology across the treatment timeline. **Row 2:** Composite multi-channel images depictin**g** the integrated distribution of proteins (green), lipids (red), and carbohydrates (blue). **Row 3-5:** Chemical heat maps illustrating the individual spatial distribution of lipid, protein, a**n**d carbohydrate, respectively. The color scales represent the relative absorbance intensity, whe**re** warmer colors denote higher concentrations. SPT: steam-pressure toasting.

However, clear heat-induced changes in relative absorbance intensity were observed for all three macronutrients. From the composite multi-channel images (Row 2) or individual protein distribution images (Row 4), the protein signal intensity decreased after SPT treatment, with the untreated control (SPT_0) showing the strongest signal in the aleurone layer and embryo regions. The lipid chemical maps (row 3) exhibited relatively stable absorption intensities during the 0-90 minutes of SPT, followed by a notable increase at 120 minutes, where the crease region, embryo and pericarp became brighter compared to the shorter processing and the untreated control. In contrast, the carbohydrate chemical maps (Row 5) displayed a reduction in the proportion of high-intensity (warm color) regions after SPT. The untreated control (SPT_0) exhibited extensive areas of high absorbance throughout the endosperm, and these areas contracted progressively as the treatment was prolonged. It should be noted that the chemical maps integrate absorbance over a narrower carbohydrate window (1178-946 cm^-1^) than the global spectra considered below (1500-900 cm^-1^); hence, the two measurements were not directly comparable.

To validate the spatial distribution patterns identified through chemical imaging, targeted spot-sampling was performed on the CDC Nasser oats harvested in 2018 and processed for 90 minutes under SPT (Figure 9), as well as on seeds harvested in 2020 and processed for 60 minutes (Figure 11). Spot spectra were extracted from anatomically distinct regions: aleurone layer, crease region, embryo, endosperm, pericarp, and sub-aleurone layer. Visible light microscopy images confirmed the morphological characteristics of each tissue. The extracted Mid-IR absorbance spectra exhibited profiles consistent with the expected biochemical composition of each tissue. The endosperm spot displayed dominant absorption in the carbohydrate fingerprint region (1500-900 cm^-1^), while the crease region and pericarp spectra were dominated by lipid-associated features, including intense CH_2_ and CH_3_ stretching bands (∼3000-2800 cm^-1^) and a prominent C=O stretching band (∼1740 cm^-1^). The aleurone layer and sub-aleurone layer spectra exhibited well-defined Amide I (∼1650 cm^-1^) and Amide II (∼1550 cm^-1^) absorption bands, and the embryo spectrum displayed a mixed profile with lipid and protein contributions. These results confirmed the chemical imaging findings and provided direct evidence that the spatial distribution of macronutrients in specific tissues of oat seeds remains unchanged after prolonged SPT, so that the spectral changes induced by the treatments reflect alterations in the molecular structure and physical state of the macronutrients rather than their spatial redistribution.

**Figure 11.**
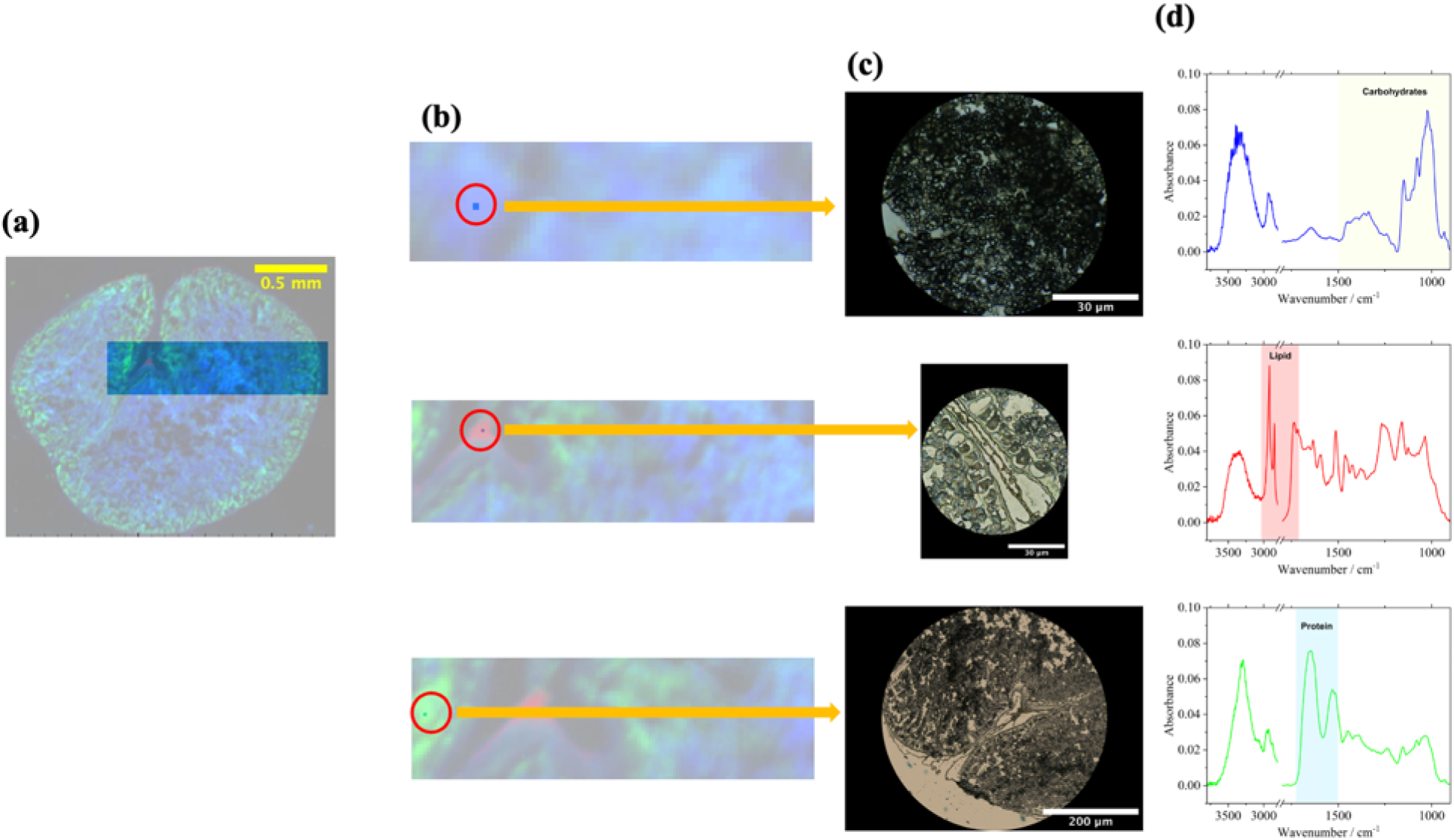
The spatial and spectral analysis of a CDC Nasser oat seed with 60 minutes of steam**­**pressure toasting harvested from Year 2020. **(a)** Mid-IR image of the intact seed cross-sectio**n,** highlighting the region of interest for targeted spot sampling. **(b)** Magnified view of the select**ed** region, with red circles indicating specific sampling spots across main anatomical structure**s:** endosperm, crease region, and sub-aleurone layer. **(c)** Visible light microscopic images corresponding to each selected spot. **(d)** Extracted Mid-IR absorbance spectra for each sp**o**t, illustrating the localized chemical profiles of carbohydrates, lipids and proteins.

As in the variety experiment, differences in overall signal intensity were observed among harvest years. The samples from 2018 and 2019 (Figure 8 and Supplemental Figure S8), prepared with an overnight hydration step, exhibited distinct spatial boundaries and strong absorption signals, whereas those from 2020 (Figure 10), prepared by an external sectioning service without the hydration step, showed reduced signal intensity, particularly in the lipid and protein maps, and a less pronounced delineation between tissue compartments. The basic tissue distribution patterns and the SPT-induced intensity trends remained largely consistent across all three years.

The global Mid-IR absorption spectra (3600-900 cm^-1^) of CDC Nasser oats across five SPT durations showed broadly similar spectral curves, in which the regions of all three major macronutrients were clearly identified, including lipids (∼3000-2800 cm^-1^ and ∼1740 cm^-1^), protein (1700-1500 cm^-1^), and carbohydrates (1500-900 cm^-1^) in Figure 12. In the carbohydrate region (1500-900 cm^-1^, panel b), the peak areas exhibited a non-monotonic and biphasic trend. The peak areas for SPT_30 and SPT_60 were smaller than that of the untreated control (SPT_0), while the peak areas for SPT_90 and SPT_120 exceeded the control. One explanation for this pattern is the sequence of the gelatinization followed by retrogradation. During the initial stage of SPT, heat and moisture disrupted the semi-crystalline structure of starch granules and the hydrogen bonds in the ordered double-helix arrangement of amylopectin, causing the granules swelling and loss of the starch crystallinity (Liu et al., 1991). After cooling, the dispersed amylose and amylopectin chains reassembled into more ordered structures, such as B- and V-type crystallites and type III resistant starch (Wang et al., 2015). All samples in this study were cooled to room temperature prior to analysis, so retrogradation could occur in every treatment group. However, longer SPT times were expected to result in more complete gelatinization of the particles and the leaching of more amylose, thereby providing more retrogradable material during the cooling phase. Consistent with previous reports, the dual autoclaving-retrogradation treatment of oat starch has been shown to increase the intensity ratio at 1047/1022 cm^-1^ and to generate type III resistant starch (Ashwar et al., 2016; Shah et al., 2016).

**Figure 12.**
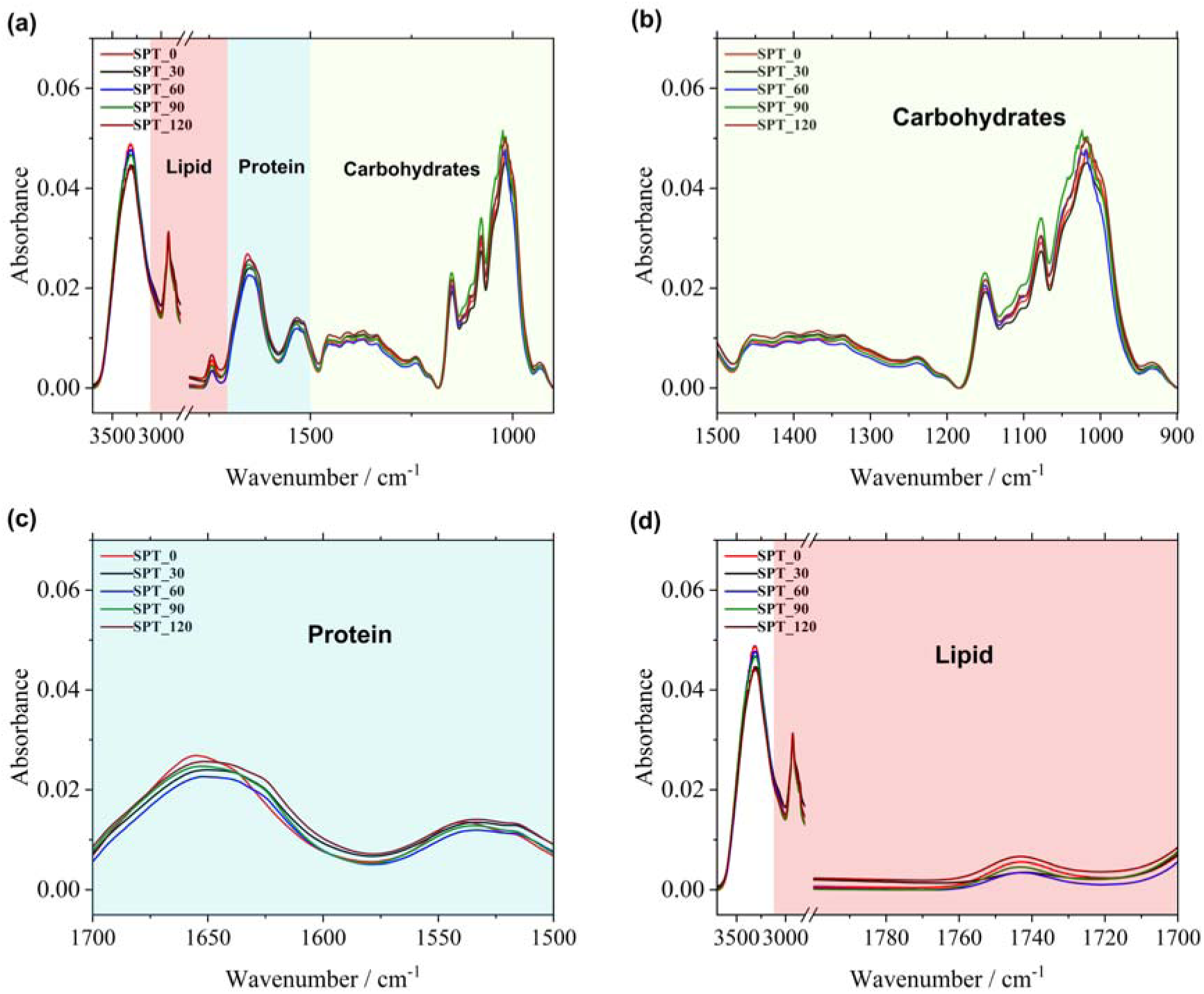
Global Mid-IR absorbance spectra of the CDC Nasser oat with different steam­pressure toasting time (0, 30, 60, 90, and 120 minutes). **(a)** The full Mid-IR spectra profiles (approx. 3600-800 cm^-1^). The spectra for the oat varieties represent the weighted average of three biological replicates and then the average of three consecutive harvest years (2018, 2019 and 2020); **(b)** Magnified view of the carbohydrate-associated region (1500-900 cm^-1^); **(c)** Magnified view of the protein-associated region (1700^-1^500 cm^-1^); **(d)** Magnified view of the lipid- associated region, illustrating spectral variations induced by the thermal processing time.

Nevertheless, this interpretation should be treated with caution, because the integrated absorbance of the carbohydrate region is mainly determined by the amount of carbohydrates within the optical path, the section thickness, and the local composition, rather than by molecular order (Baker et al., 2014). In addition, the spectral bands in this region overlap significantly, and the gelatinization caused the intensity to redistribute among them, rather than eliminating the C­O, C-C, and C-O-C vibrational modes that contribute to the absorbance. For example, the band corresponding to the ordered structure near 1047 cm^-1^ flattened, the amorphous band near 1022 cm^-1^ increased, and the band near 995 cm^-1^, which was likewise associated with ordered structure, decreased (Xu et al., 2021). Therefore, a decrease in the integral was not an inevitable result of amorphization, and the relationship between the ordered structure of starch and the infrared peak ratio was not a simple linear one (Warren et al., 2016). The absorbance of proteins decreased within the same treatment range, and the Maillard reaction products formed between reducing sugars and amino acids absorbed in the same fingerprint region: infrared characterization of glucose-alanine melanoidins assigned a strong C-O band at 1080-1035 cm^-1^ and a C-C band of the retained carbohydrate skeleton near 920 cm^-1^, which contribute to the same broad absorption region (Mohsin et al., 2022). Overall, the biphasic trend reported in our study is a spectroscopic observation and is consistent with the pattern of gelatinization-retrogradation. Further confirmation in subsequent experiments will be required, which will need to explicitly evaluate the short-range ratio, X-ray diffraction, differential scanning calorimetry or resistant starch analysis.

In the protein region (1700-1500 cm^-1^, panel c), the peak centers of both Amide I and II bands in all treated samples shifted to lower wavenumbers and both bands broadened, while the Amide I peak increased slightly in height and the Amide II peak height remained essentially unchanged. This spectral shift reflects a transition in the composition of the protein secondary structure from high wavenumber α-helices (∼1653 cm^-1^) and random coils (∼1645 cm^-1^) to low wavenumber β - sheets (∼1636 and 1614 cm^-1^). Similar literature indicated that the intensity of Amide I shifted toward the 1630-1613 cm^-1^ region during the heating of wheat gluten, which was attributed to the formation of β-sheets (Georget and Belton, 2006). The simultaneous broadening of these two spectral bands was consistent with this transition, as the predominantly disordered group transformed into several overlapping β-sheets and residual components. This broadened and diversified the distribution of the hydrogen-bonding environment contributing to the amide band. A slight increase in the intensity of Amide I band did not require an increase in protein content. This was because the intermolecular β-sheet structure gave rise to a strong low-wavenumber Amide I band (∼1620 cm^-1^), whose intensity stemmed from transition dipole coupling between the C=O oscillators of the aligned peptides (Barth, 2007). Therefore, the growth of this strong absorption component increased the apparent intensity of the Amide I band as the aggregation proceeded. The intensities of Amide I and Amide II did not decrease because of processing, and the global spectrum showed no loss of amide-active proteins. Structural rearrangement was manifested by changes in band positions and widths without any decrease in intensity. The corresponding secondary-structure proportions are quantified by band deconvolution in Section 3.2.2.

In the lipid region (∼3000-2800 cm^-1^ and ∼1740 cm^-1^, panel d), the intensities of CH_2_/CH_3_ and the C=O stretching bands remained at or slightly below the levels of the untreated control from 30 to 90 minutes of SPT but exceeded the control group at 120 minutes. This response indicated that a SPT treatment longer than 90 minutes was required to induce substantial changes in the physical state and lipid accessibility within the seed tissue. In native oat seeds, most of the lipid is stored in oil bodies, in which triacylglycerol droplets are embedded within a phospholipid monolayer surrounding oleosin proteins. This structure is highly conserved among seeds, providing physical stability to the droplets (Tzen et al., 1993). During intermediate toasting (30­90 minutes), the surface of oil bodies may become partially unstable. Nonetheless, this lipid surface may still retain its structural integrity, keeping lipids in a bound state; thus, its infrared absorption characteristics remain largely unchanged. In contrast, during the 120 minutes SPT, the cumulative thermal damage may severely disrupt the oleosin-phospholipid interface and released the bound lipids into a free state. The increased thermal mobility of these released lipids facilitates their migration into the intercellular space, thereby increasing the spatial coverage and local concentration of lipids in the region. In addition, prolonged thermal exposure may promote lipid oxidation, leading to the formation of secondary oxidation products (such as aldehydes and ketones). Former research reported that secondary carbonyl products generated by the thermal oxidation of edible oils were absorbed near the ester C=O bands at approximately 1740 cm^-1^ and in the 1700-1730 cm^-1^ region (Guilléή and Cabo, 2002). These previous outcomes overlap with the lipid integration window used in our study (1770-1719 cm^-1^). Both lipid redistribution and the accumulation of oxidation products might increase the total absorbance in this window, and the current data do not allow for distinguishing between the two.

#### 3.2.2 Changes in protein secondary structures induced by SPT

To further investigate protein secondary structural changes induced by SPT, peak-fitting deconvolution of the Amide I region (1700-1600 cm^-1^) was conducted. Deconvolution results for CDC Nasser oats harvested in 2018 were presented in Figure 13, with results from 2019 and 2020 provided in Supplemental Figure S11. In the untreated control (SPT_0, panel a), the *a* - helix subpeak (∼1653 cm^-1^, pink peak) was relatively small, and the random coil component (∼1645 cm^-1^, green peak) was clearly visible as a distinct subpeak. After SPT with different durations, *a* -helix subpeaks markedly increased in amplitude across all treated samples (panels b-e), while the random coil sub-peaks diminished substantially and became largely obscured by the enlarged a -helix peak in treated samples. Additionally, the 0-sheet sub-peaks (∼1636 and ∼1614 cm^-1^, purple and cyan) also exhibited a notable increase in amplitude after thermal treatment, with the most substantial increases observed in the SPT_90 and SPT_120 samples. The *β*-turn subpeaks (∼1684 and ∼1661 cm^−1^, orange and yellow) remained relatively consistent across treatments. These deconvolution results provided clear visual evidence that STP induced redistribution of protein secondary structures, characterized by an increase in *β*-sheet content at the expense of random coil structures.

**Figure 13.**
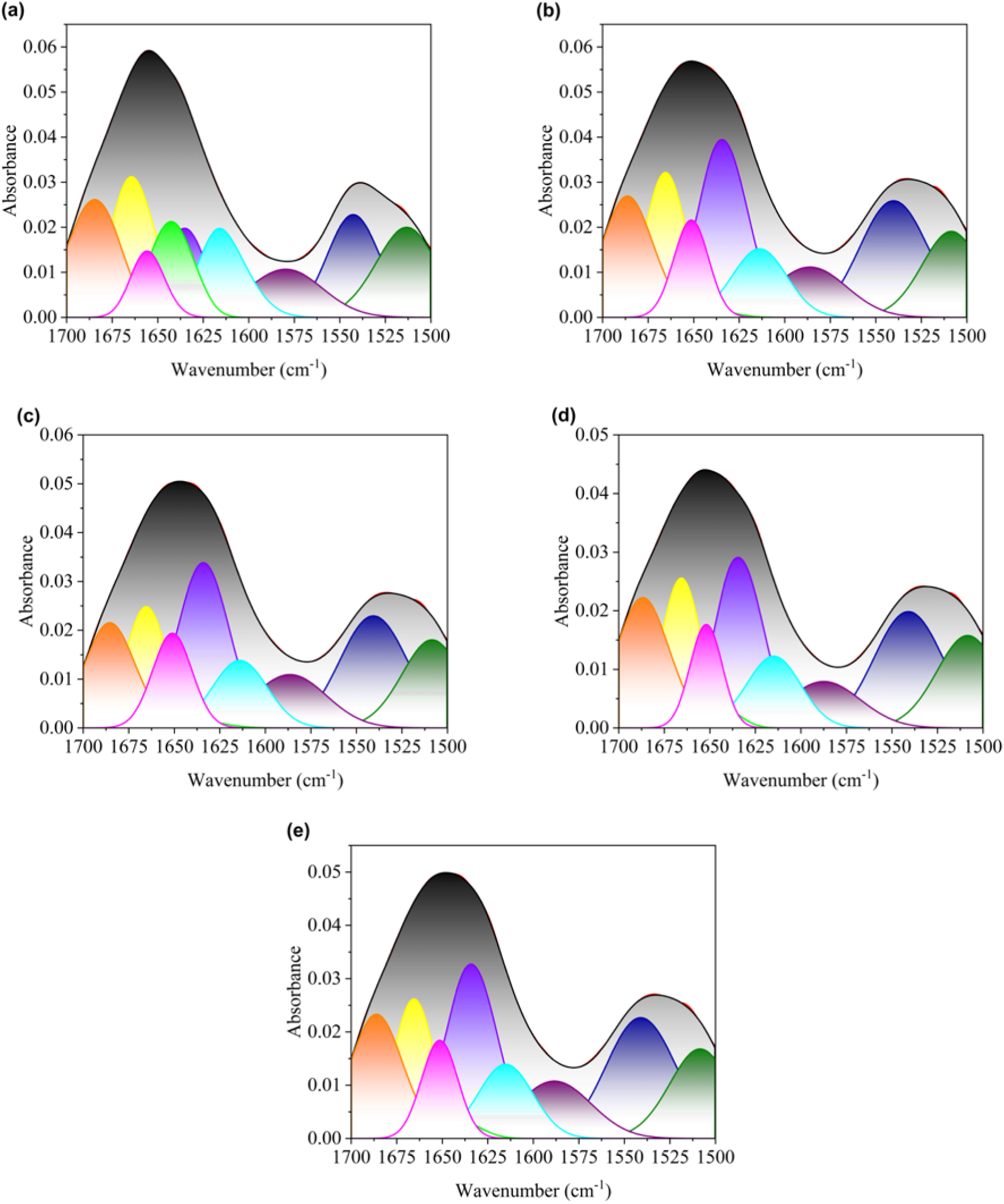
Peak-fitting deconvolution of the Mid-IR Amide I band (1700-1600cm^-1^) tracking secondary protein structural changes in CDC Nasser oats harvested in 2018 during steam- pressure toasting. The panels represent the heating durations: **(a)** 0 min (control), **(b)** 30 min, **(c)** 60 min, **(d)** 90 min, and **(e)** 120 min. The black solid line represents the original raw spectrum, and the red solid line indicates the overall fitted curve. The color-filled subpeaks correspond to protein secondary structures: 0-turns (orange and yellow peaks, 1700-1660 cm^-1^), a -helices (pink peak, 1660-1650 cm^-1^), random coils (green peak, 1650-1640 cm^-1^), -sheets (purple and cyan peaks, 1640-1600 cm^-1^).

Quantitative analysis of protein secondary structure proportions across the five SPT durations (Figure 14) confirmed statistically significant treatment effects for *β*-sheet (*P = 0.003*) and random coil (*P = 0.026*), while *a*-helix (*P = 0.418*), *β*-turn (*P = 0.239*), and the *a*-helix to *β* - sheet ratio (*P = 0.811*) showed no significant differences. Multi-treatment comparisons provided clear pairwise separations for both significant parameters. For *β*-sheet, the untreated control group (SPT_0, group “a”) was significantly lower than all four heat treated groups (SPT_30, SPT_60, SPT_90, and SPT_120, all group “b”), indicating that 30 minutes of SPT significantly increases the *β*-sheet contents but no further significant changes beyond 30 min. Conversely, SPT_0 (group “a”) exhibits a significantly higher proportion of random coils than all treated groups (SPT_30, SPT_60, SPT_90, and SPT_120, all group “b”), confirming that 30 minutes of SPT was sufficient to significantly reduce the disordered random coil content. The increase in *β* - sheets and the decrease in random coils at 30 minutes of SPT strongly suggest a direct structural transition from a disordered random coil to an ordered *β*-sheet during heat treatment. Compared to SPT_0, the relative proportion of *a*-helices increased in all SPT treatment while the difference was not statistically significant (*P = 0.418*), although the subpeaks of a-helix in the deconvoluted spectra was visually evident (Fig. 13). Under all treatment conditions, the 0-turn and the ratio of α-helix to *β*-sheet remained stable, which indicated that the primary effect of SPT was a redistribution of conformations from disordered random coils to ordered *β*-sheet.

**Figure 14.**
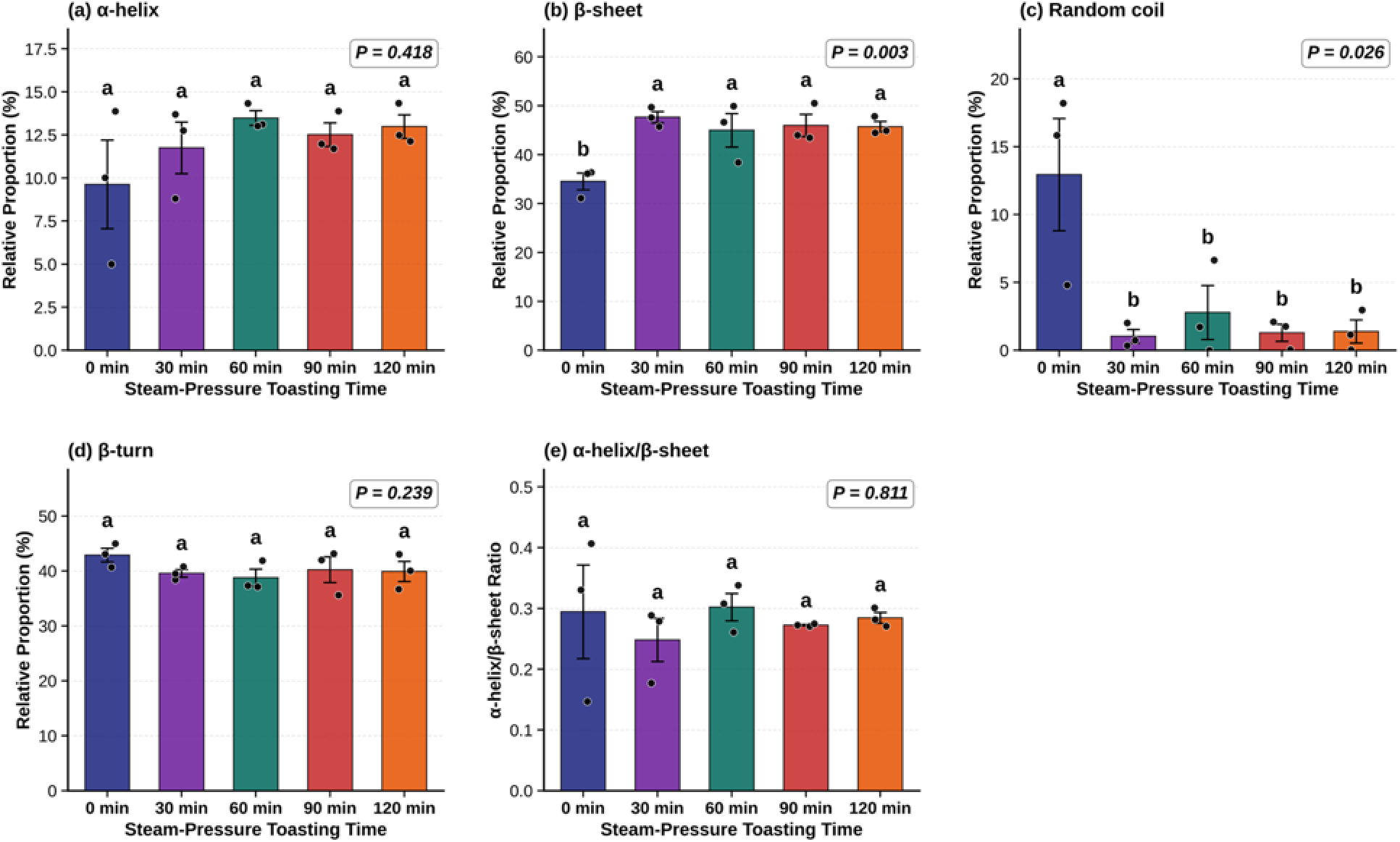
Relative proportions of secondary protein structures in CDC Nasser oats acro**s**s different steam-pressure toasting durations (0, 30, 60, 90, and 120 minutes). Quantitative resu**lt**s were gained from the Mid-IR Amide I band deconvolution. Panels display the calculat**ed** percentage for **(a)** -helix, **(b)** -sheet, **(c)** random coil, **(d)** -turn, and **(e)** the ratio of -hel**i**x to -sheet. For each treatment within each harvest year, three seed cross-sections (sub-sampl**es**) were measured and averaged to obtain a single block-level observation. Bar heights indic**ate** mean value of three harvest years, and the individual black dots represent the block-level mea**n**s from 2018, 2019, and 2020 (n = 3). Error bars are the standard error means (SEM). ANOV**A** analysis was performed under randomized complete block design (RCBD) with harvesting ye**ar** as the blocking factor. Overall model *P*-values are displayed in the upper right corner of eac**h** panel. Means with the different letters are significant different (P < 0.05), and multi-treatme**nt** comparison is LSD method.

During hydrothermal processing, the thermal energy broke the weak hydrogen bonds and hydrophobic interactions that stabilize the native fold. Driven by hydrophobic interactions and thermodynamic stability, the unfolded peptide chains reassemble into highly ordered and compact intermolecular *β*-sheet aggregates held together by strong intermolecular hydrogen bonds. Ma and Harwalkar (1988) demonstrated that preheating oat globulin gradually reduced the enthalpy of denaturation using differential scanning calorimetry while increasing the denaturation temperature and narrowing the endothermic peak. This suggested that the conformation of the preheated proteins became more compact and correlated, exhibiting greater thermal stability and synergy. Their salt-dependence data also showed that hydrophobic interactions contributed to this stability. In other studies, the same transition has also been reported in plant proteins using spectroscopy. For example, heating wheat gluten caused the low-frequency Amide I peak to shift from approximately 1630 to 1613 cm^-1^, which could be attributed to *β*-sheets (Georget and Belton, 2006). Legume proteins increased intermolecular *β* - sheet aggregation at the expense of disordered conformations after heat treatment (Carbonaro et al., 2012). This transition from disordered random coils to ordered *β*-sheet structures is the sign of heat induced denaturation and aggregation of cereal proteins. This suggests that the structural transition from the disordered to the ordered conformation occurs rapidly under SPT with 30 minutes, and extending the treatment time does not further alter the equilibrium between these two structural components. One plausible explanation was that the thermally unstable and solvent-accessible disordered structures were largely exhausted early in the process, so the changes that occurred later were primarily due to chemical modifications, such as Maillard reactions and lipid oxidation, rather than further conformation rearrangements.

The decrease in protein intensity observed in the FTIR chemical maps (Figure 8) did not disagree with the increase in the 0-sheet fraction. It should be interpreted in conjunction with the global spectrum (Figure 12), in which the Amide I intensity did not decrease (Section 3.2.1). These two intensity measurements were fundamentally different. The map integrated the Amide I absorbance across the entire cross-sections using a globar-source FTIR instrument, whereas the point spectrum sampled the aleurone layer and endosperm using a synchrotron source. The map intensity was also sensitive to the slice thickness and local tissue density. Neither of these measurements indicated a loss of protein. Aggregation reduced the extractable protein content of heat-treated oats from approximately 75% to 36%, but this represented a loss of extractability, not a loss of protein. The aggregated proteins remained within the kernel and retained their amide activity (Runyon et al., 2015). The Maillard reaction proceeded in parallel, reducing the availability of active lysine and proteins. Since it modified the side-chain amino group rather than the main chain C=O bond, its direct contribution to amide strength was expected to be limited (Pahm et al., 2008). The spectrum clearly showed changes in band position, and the shift of the Amide I peak center toward lower wavenumber was an observation of band shape and was not related to absolute intensity. However, this provided direct spectroscopic evidence of a transition from the random coil at high wavenumbers (∼1645 cm^-1^) to the *β*-sheet at lower wavenumbers, which was consistent with our deconvolution results described above (∼1636 and ∼1614 cm^-1^) (Barth, 2007; Georget and Belton, 2006). These structural changes are significant for protein function and nutritional quality. Carbonaro et al. (2012) reported a strong negative correlation between the relative content of *β*-sheet structures and the in vitro protein digestibility of legumes, grains, and animal foods. They proposed that intermolecular *β*-sheet aggregates formed during heating treatment were the primary factor affecting digestibility, based on the premise that tightly packed hydrogen-bonded sheets restrict the access of proteolytic enzymes to cleavage sites. A similar correlation was observed between a high *β*-sheet ratio and reduced protein availability using synchrotron-based infrared spectroscopy to identify protein sources in feed (Yu et al., 2004b). However, in the specific background of STP of ruminant feed, a shift toward slower protein degradation may be beneficial, since STP was intended to reduce ruminal degradation and shifted the supply of amino acids to the small intestine (Goelema et al., 1998; Yu et al., 2000). Therefore, the optimal heating duration was determined by a balance between the reduced rate of ruminal degradation and the risk of reduced intestinal digestibility (van der Poel et al., 2005). The results of this study were also related to that balance: the *β*-sheet transformation was completed after 30 minutes, while Maillard and lipid-related changes continued to accumulate up to 120 minutes, suggesting that a shorter SPT duration could capture most of the desired structural modifications while limiting further thermal damage. Since the current study characterized only molecular structures, this inference needs to be further confirmed through in situ or in vitro degradation and digestibility tests, as was done for steam-pressure-toasting faba beans (Rodríguez Espinosa and Yu, 2025).

## 4. Conclusion

This study utilized synchrotron-based mid-infrared spectroscopy and FTIR chemical imaging to investigate the distribution of macronutrients and the effects of variety and steam-pressure toasting (SPT) duration on protein secondary structure in cool-season oats. Three main conclusions can be drawn from this study.

Firstly, the spatial distribution of macronutrients was a conserved structural feature of oat seeds, such as carbohydrates in the endosperm, proteins in the aleurone/sub-aleurone layer and embryo, and lipids in the crease region and embryo. This pattern was consistent across all four varieties and remained unchanged after SPT, suggesting that processing alters the molecular state of the macronutrients rather than their location at the tissue level.

Secondly, oat varieties had only a minor effect on protein secondary structure. Against a common main-chain conformation dominated by 0-sheet, the variety effects on the a-helix, *β* - turn, and ratio of *a*-helix to *β*-sheet were different. SPT fundamentally reorganized the protein matrix, driving a transition from disordered random coils to ordered *β*-sheet aggregates. The significant increase in *β*-sheets and decrease in random coils were already complete after 30 minutes, with no further significant changes observed up to 120 minutes. This transition was rapid and consistent with heat-induced denaturation and aggregation of oat globulin. Importantly, there was no contradiction between the reduction in protein signals observed in the chemical map and the increase in the proportion of *β*-sheets: the global spectra showed that the intensity of the Amide I band has not decreased. Therefore, there was no implied loss of protein, and the aggregated protein remained within the kernel while retaining its amide activity. Furthermore, the increase in the *β*-sheet proportion was related to the position and shape of the Amide I band, not to its absolute intensity.

Thirdly, the carbohydrate matrix presented a biphasic response to SPT: starch gelatinization and a loss of crystallinity occurred during the first 30-60 minutes, followed by retrogradation and the formation of new ordered and resistant-starch structures between 90 and 120 minutes, accompanied by increasing Maillard and lipid-oxidation contributions. Overall, these findings suggested that oat varieties had a relatively minor effect on the protein secondary structures of oats, but SPT treatment reorganized the protein matrix from a disordered state into an ordered aggregated conformation, while simultaneously restructuring the starch and lipids. Since aggregated proteins rich in *β*-sheets were typically less soluble and less readily digested, and since most of the structural reorganization occurred within the first 30 minutes while Maillard and lipid-related changes accumulated over a longer period, the results suggested that a shorter SPT duration may achieve the desired structural modifications while limiting the associated nutritional losses. Because this study only characterized the molecular structure, this inference needs to be confirmed through direct digestibility measurements. These structural-function relationships bear directly on the protein digestibility, solubility, and overall nutritional functionality of processed oats, and provide a spectroscopic basis for optimizing thermal processing conditions.

## Declaration of competing interest

The authors declare that they have no known competing financial interests or personal relationships that could have appeared to influence the work reported in this paper.

## Acknowledgements

We acknowledge Dr. Aaron Beattie of Crop Development Center (CDC) at the University of Saskatchewan to provide us the seeds for this research. The research described in this paper was performed at the Canadian Light Source Inc. (CLS), a national research facility of the University of Saskatchewan, which is supported by the Canada Foundation for Innovation (CFI), Natural Science and Engineering Research Council (NSERC), National Research Council (NRC), Canadian Institutes of Health Research (CIHR), Government of Saskatchewan and University of Saskatchewan. The Ministry of Agriculture Strategic Research Chair Programs fund from the Saskatchewan Pulse Growers (SPG), the Natural Sciences and Engineering Research Council of Canada (NSERC-Individual Discovery Grant and NSERC-CRD Grant), SaskCanola, the Saskatchewan Agriculture Strategic Research Chair Program Fund, the Agricultural Development Fund (ADF), SaskMilk, the Saskatchewan Forage Network (SNK), and the Western Grain Research Foundation (WGRF) are acknowledged. Ganqi Deng is financially supported by the China Scholarship Council (CSC) in the University of Saskatchewan, Canada.

## Declaration of generative AI and AI-assisted technologies in the writing process

During the preparation of this work the author used ChatGPT-5.6 (OpenAI) and Claude-5 (Anthropic) to refine the language and improve the coherence and clarity of the manuscript. After using this tool/service, the author reviewed and edited the content as needed and takes full responsibility for the content of the publication.

## Notes

### Competing Interest Statement

The authors have declared no competing interest.

