## Supplemental Material for "Macronutrient Distribution and Protein Secondary Structure of Cool-Season Oats Revealed by Synchrotron-Based Mid-IR Spectroscopy and FTIR Chemical Imaging: Effects of Variety and Steam-Pressure Toasting Duration"

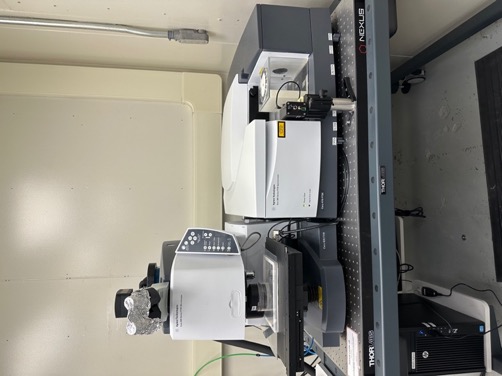

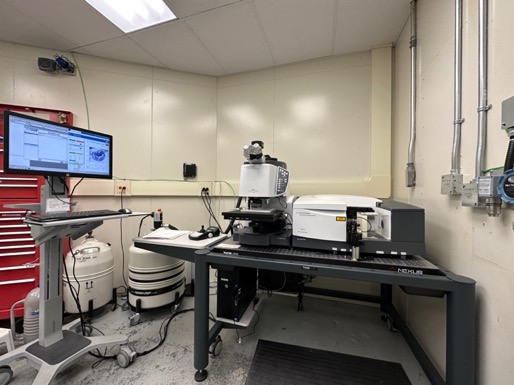

(a)	(b)

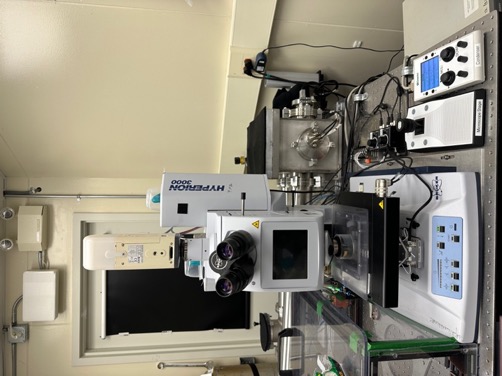

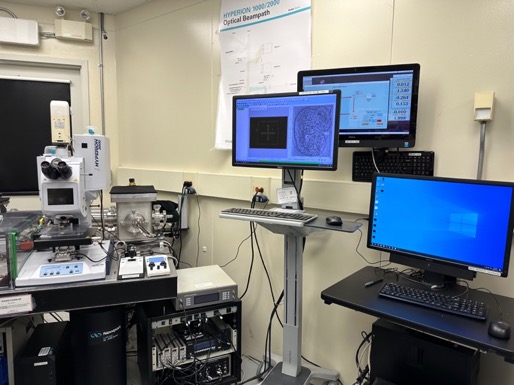

(c)	(d)

**Figure S1.** Mid-infrared spectrometers at the Mid-IR beamline of the Canadian Light Source (Saskatoon, Canada). (a, b) The Hyperion 3000 FPA FTIR microscope, equipped with a globar internal thermal light source, is used for Transmission Imaging to map the spatial distribution of protein, carbohydrates, and lipids in oat seed cross-sections. (c, d) The Hyperion 3000 MCT infrared microscope, coupled to the synchrotron light source, is used for high-resolution single-point spectral acquisition of oat seed sections.

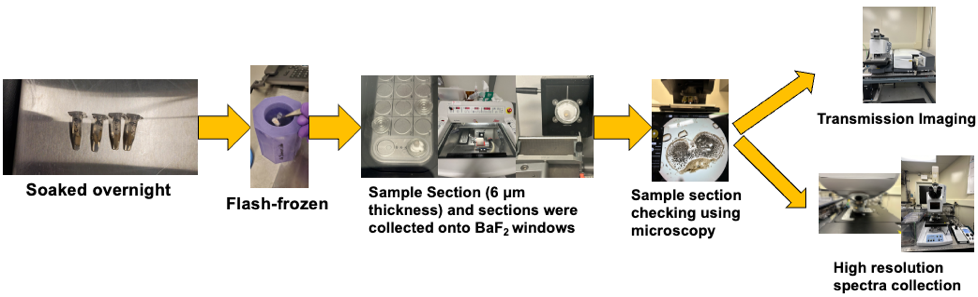

**Figure S2.** Workflow for sample preparation and Mid-IR data collection. Oat seeds were soaked overnight, rapidly frozen in liquid nitrogen, and sectioned at a thickness of 6 μm using a cryostat. The sections were collected onto BaF2 windows and visually inspected under a light microscope to verify section quality and tissue integrity. The prepared sections were then analyzed using two complementary techniques: (1) Transmission chemical imaging using the Hyperion 3000 FPA infrared microscope with a globar internal thermal source, and (2) the Hyperion 3000 MCT infrared microscope coupled with synchrotron light source.

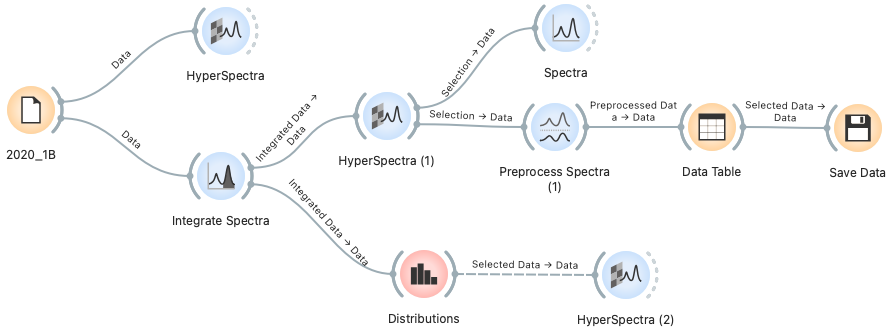

**Figure S3.** Example of the data analysis workflow for transmission imaging data using Quasar software (Orange-Quasar). Raw hyperspectral data (e.g., 2020_1B) was loaded and visualized using the HyperSpectra widget. The Integrate Spectra widget was used to determine the integrated absorbance regions corresponding to protein, carbohydrates and lipids. The distribution widget was then used to define intensity thresholds for the RGB color channels (red, green, and blue upper limits), which were applied in the HyperSpectra (1) widget to generate a composite RGB chemical distribution map as well as separate heatmaps for each biochemical component.

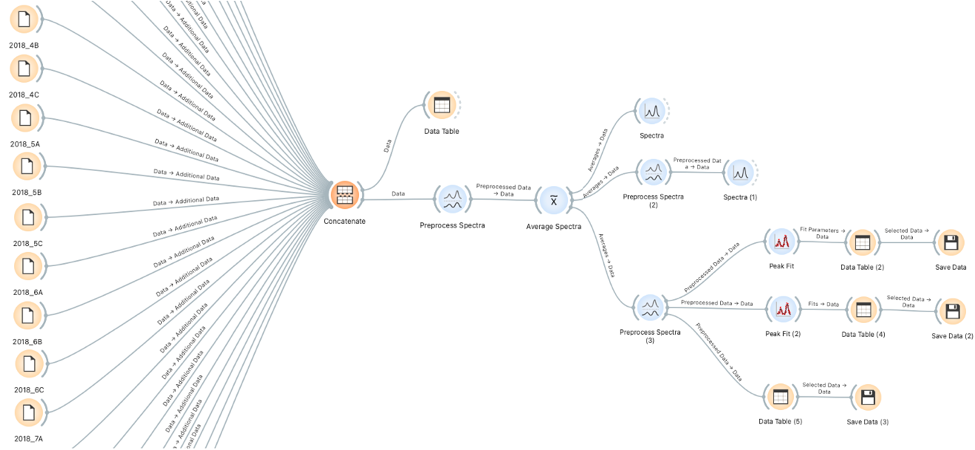

**Figure S4.** The example of high-resolution spectral data analysis and peak-fitting workflow in Quasar software (shown for samples harvested in 2018). The Concatenate widget was used to combine individual spectral files from each oat varieties and steam-pressure toasting durations with replicates from 2018. The concatenated spectra were preprocessed (cut, Gaussian smoothing, baseline correction, and normalization), and then averaged using the Average Spectra widget by source ID to produce a representative spectrum for each sample. The averaged spectra were directed through two parallel branches: i) Preprocess Spectra (2) widget applied a Savitzky–Golay second-derivative filter to verify the peak positions and visualized by the Spectra (1) widget; and ii) Preprocess Spectra (3) cut the spectra to the Amide I/II region (1700-1500 cm^-1^) for peak-fitting deconvolution using the Peak Fit and Peak Fit (2) widgets. Then fitted parameters and fitted spectra were exported as CSV files by the Save Data widgets for subsequent analysis and visualization in Origin software.

**Replicate 2:**

CDC Haymaker	Summit	CDC Nasser	CDC Arborg

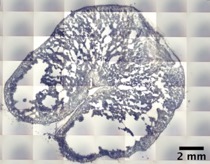

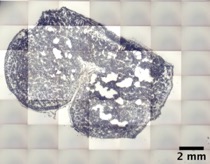

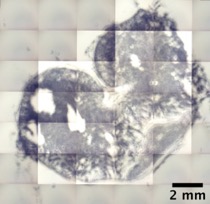

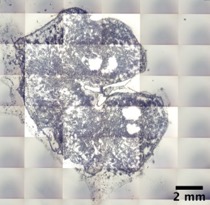

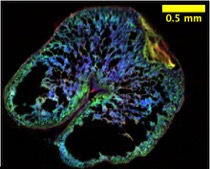

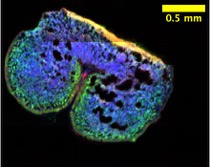

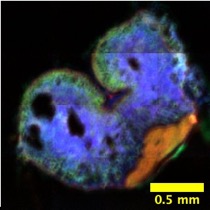

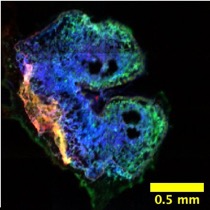

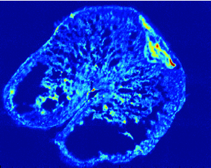

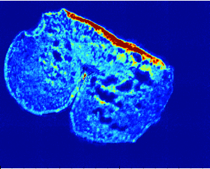

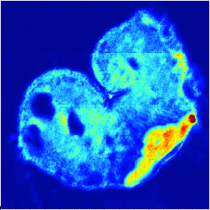

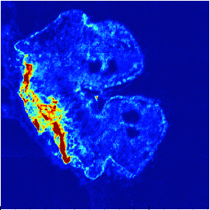

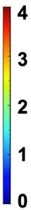

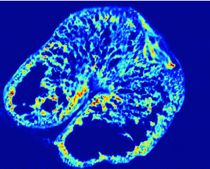

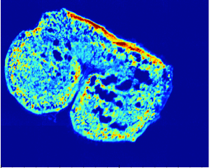

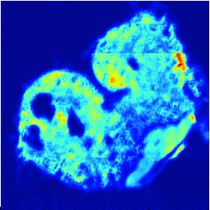

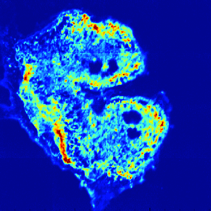

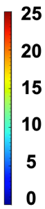

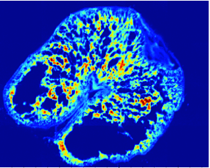

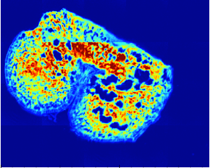

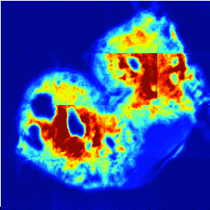

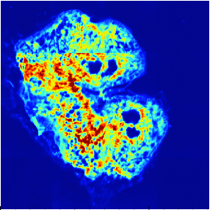

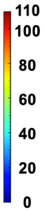

**Replicate 3:**

CDC Haymaker	Summit	CDC Nasser	CDC Arborg

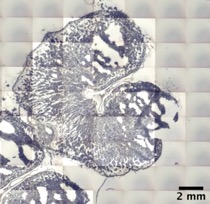

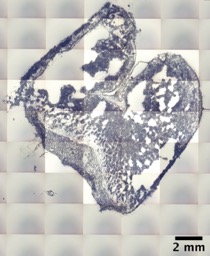

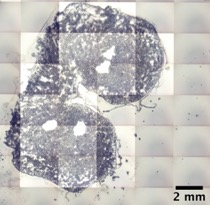

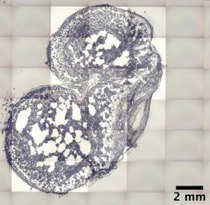

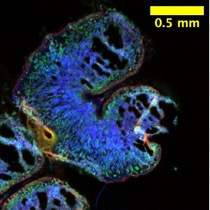

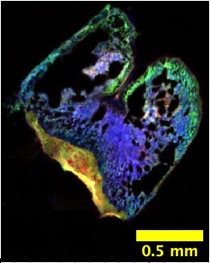

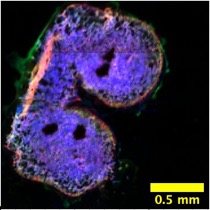

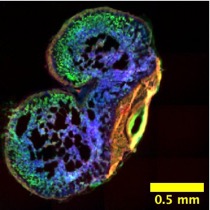

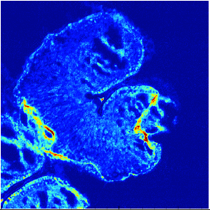

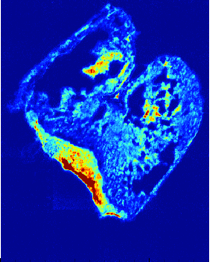

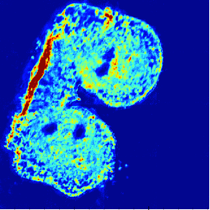

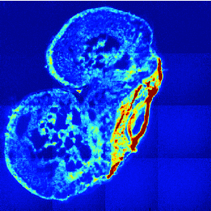

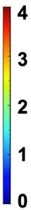

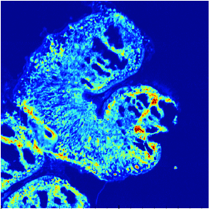

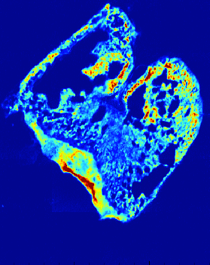

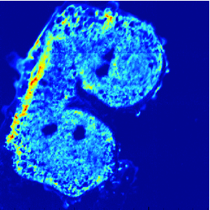

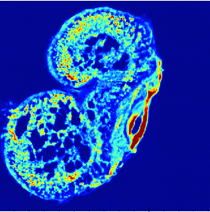

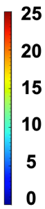

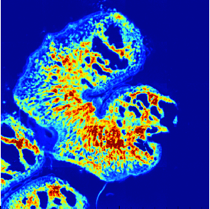

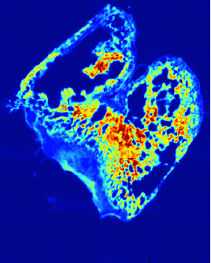

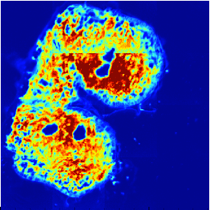

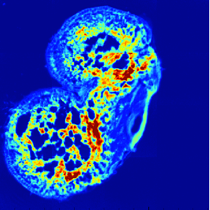

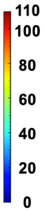

**Figure S5.** Second and third replicates for the spatial distribution of macronutrients (lipids, protein, carbohydrates) within the seed cross-sections of four cool-season oat varieties (CDC Haymaker, Summit, CDC Nasser, CDC Arborg) harvested from Year 2018. **Row 1:** Visible light microscopy images displaying the intact seed morphology. **Row 2:** Composite multi-channel images illustrating the combined distribution of protein (green), carbohydrate (blue) and lipid (red). **Rows 3-5:** Individual chemical heat maps for the lipids, protein and carbohydrates distributions, respectively. The color scales represent the relative absorbance intensity, where warmer colors denote higher concentrations.

**Replicate 2:**

CDC Haymaker	Summit	CDC Nasser	CDC Arborg

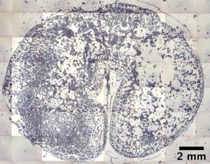

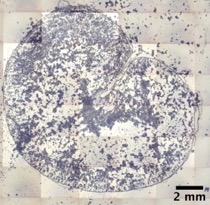

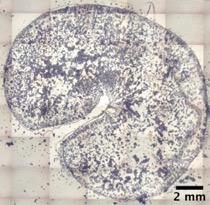

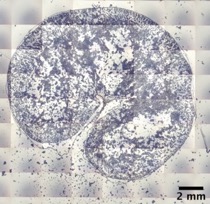

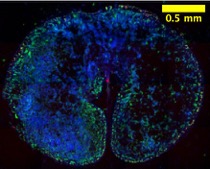

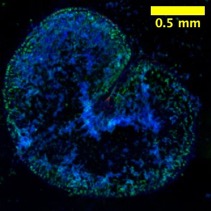

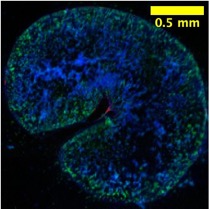

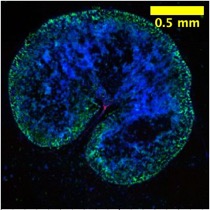

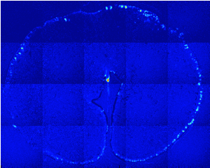

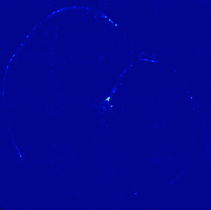

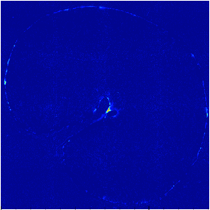

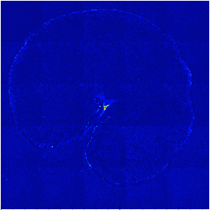

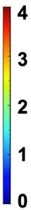

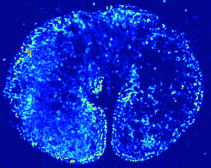

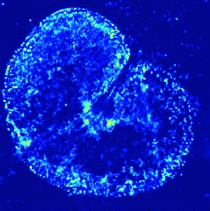

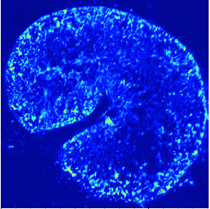

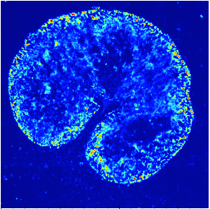

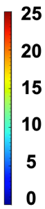

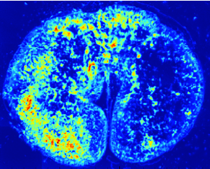

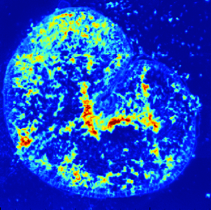

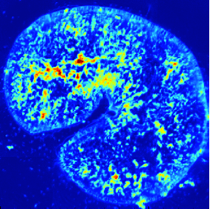

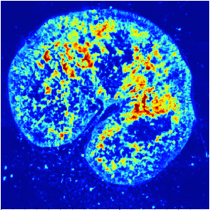

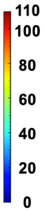

**Replicate 3:**

CDC Haymaker	Summit	CDC Nasser	CDC Arborg

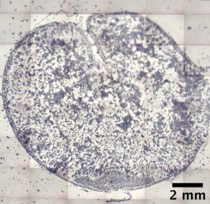

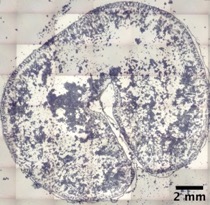

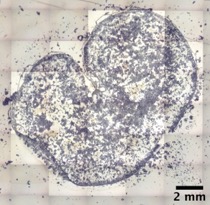

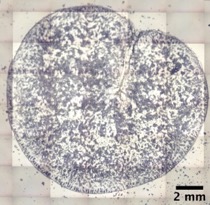

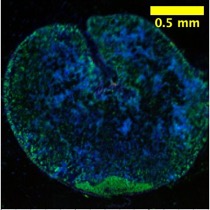

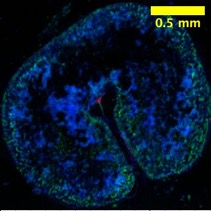

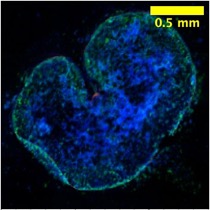

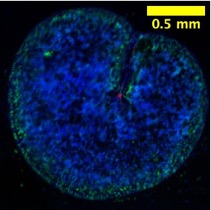

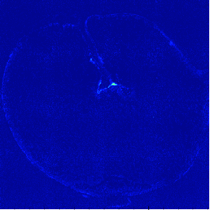

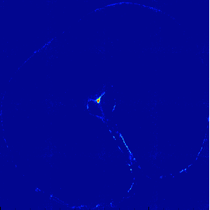

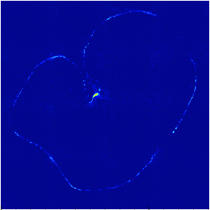

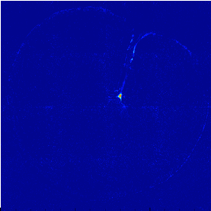

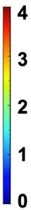

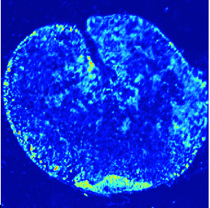

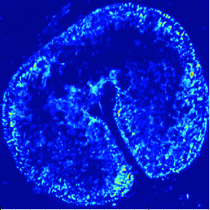

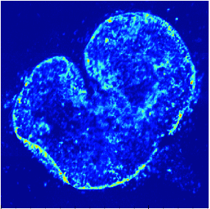

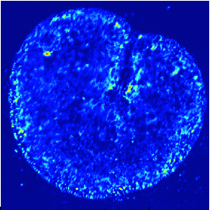

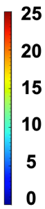

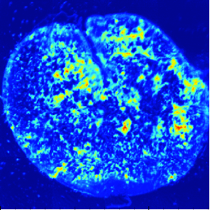

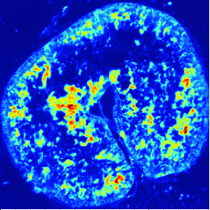

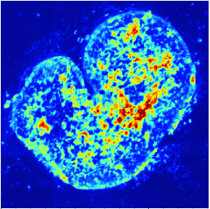

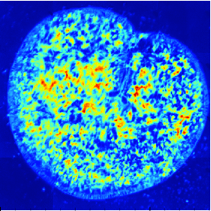

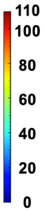

**Figure S6.** Second and third replicates for the spatial distribution of macronutrients (lipids, protein, carbohydrates) within the seed cross-sections of four cool-season oat varieties (CDC Haymaker, Summit, CDC Nasser, CDC Arborg) harvested from Year 2020. **Row 1:** Visible light microscopy images displaying the intact seed morphology. **Row 2:** Composite multi-channel images illustrating the combined distribution of protein (green), carbohydrate (blue) and lipid (red). **Rows 3-5:** Individual chemical heat maps for the lipids, protein and carbohydrates distributions, respectively. The color scales represent the relative absorbance intensity, where warmer colors denote higher concentrations.

**Replicate 2:**

SPT_0	SPT_30	SPT_60	SPT_90	SPT_120

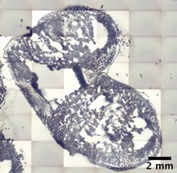

**Replicate 3:**

SPT_0	SPT_30	SPT_60	SPT_90	SPT_120

**Figure S7.** Second and third replicates for the spatial distribution of macronutrients within the seed cross-sections of the CDC Nasser oat with different durations of steam-pressure toasting (0, 30, 60, 90, and 120 minutes) harvested from Year 2020. **Row 1:** Visible light microscopy images showing the intact seed morphology across the treatment timeline. **Row 2:** Composite multi-channel images depicting the integrated distribution of proteins (green), lipids (red), and carbohydrates (blue). **Row 3-5:** Chemical heat maps illustrating the individual spatial distribution of lipid, protein, and carbohydrate, respectively. The color scales represent the relative absorbance intensity, where warmer colors denote higher concentrations. SPT: steam-pressure toasting.

**Replicate 1:**

SPT_0	SPT_60	SPT_90	SPT_120

**Replicate 2:**

SPT_0	SPT_60	SPT_90	SPT_120

**Replicate 3:**

SPT_0	SPT_60	SPT_90	SPT_120

**Figure S8.** Spatial distribution of macronutrients within the seed cross-sections of the CDC Nasser oat with different durations of steam-pressure toasting (0, 60, 90, and 120 minutes) harvested from Year 2019 including three replicates. **Row 1:** Visible light microscopy images showing the intact seed morphology across the treatment timeline. **Row 2:** Composite multi-channel images depicting the integrated distribution of proteins (green), lipids (red), and carbohydrates (blue). **Row 3-5:** Chemical heat maps illustrating the individual spatial distribution of lipid, protein, and carbohydrate, respectively. The color scales represent the relative absorbance intensity, where warmer colors denote higher concentrations. SPT: steam-pressure toasting.

**Replicate 2:**

SPT_0	SPT_30	SPT_60	SPT_90	SPT_120

**Replicate 3:**

SPT_0	SPT_30	SPT_60	SPT_90	SPT_120

**Figure S9.** Second and third for the spatial distribution of macronutrients within the seed cross-sections of the CDC Nasser oat with different durations of steam-pressure toasting (0, 30, 60, 90, and 120 minutes) harvested from Year 2020. **Row 1:** Visible light microscopy images showing the intact seed morphology across the treatment timeline. **Row 2:** Composite multi-channel images depicting the integrated distribution of proteins (green), lipids (red), and carbohydrates (blue). **Row 3-5:** Chemical heat maps illustrating the individual spatial distribution of lipid, protein, and carbohydrate, respectively. The color scales represent the relative absorbance intensity, where warmer colors denote higher concentrations. SPT: steam-pressure toasting.

**Year 2019:**

**Year 2020:**

**Figure S10.** Peak-fitting deconvolution of the Mid-IR Amide I region (1700-1600 cm^-1^) for determining the protein secondary structures of four cool-season oat varieties harvested in Year 2019 and 2020. The panels stand for **(a)** CDC Haymaker, **(b)** Summit, **(c)** CDC Nasser, and **(d)** CDC Arborg. In each panel, the black solid line represents the original raw spectrum, and the red solid line indicates the overall fitted curve. The color-filled subpeaks under the curve correspond to protein secondary structures based on specific wavelength ranges: 𝛽-turn structures (1700-1660 cm^-1^) are represented by the orange (~1684 cm^-1^) and yellow (~1661 cm^-1^) peaks; the 𝛼-helix structure (1660-1650 cm^-1^) is represented by the pink (~1653 cm^-1^) peak; the random coil (1650-1640 cm^-1^) is represented by the green (~1645 cm^-1^) peak; and 𝛽-sheet structures (1640-1600 cm^-1^) are represented by the purple (~1636 cm^-1^) and cyan (~1614 cm^-1^) peaks.

**Year 2019:**

**Year 2020:**

**Figure S11.** Peak-fitting deconvolution of the Mid-IR Amide I band (1700-1600 cm^-1^) tracking secondary protein structural changes in CDC Nasser oats harvested in Year 2019 and 2020 during steam-pressure toasting. The panels represent the heating durations: **(a)** 0 min (control), **(b)** 30 min, **(c)** 60 min, **(d)** 90 min, and **(e)** 120 min. The black solid line represents the original raw spectrum, and the red solid line indicates the overall fitted curve. The color-filled subpeaks correspond to protein secondary structures: 𝛽-turns (orange and yellow peaks, 1700-1660 cm^-1^), 𝛼-helices (pink peak, 1660-1650 cm^-1^), random coils (green peak, 1650-1640 cm^-1^), and 𝛽-sheets (purple and cyan peaks, 1640-1600 cm^-1^).
